# Rewritable Two-Dimensional DNA-Based Data Storage with Machine Learning Reconstruction

**DOI:** 10.1101/2021.02.22.432304

**Authors:** Chao Pan, S. Kasra Tabatabaei, SM Hossein Tabatabaei Yazdi, Alvaro G. Hernandez, Charles M. Schroeder, Olgica Milenkovic

## Abstract

DNA-based data storage platforms traditionally encode information only in the nucleotide sequence of the molecule. Here we report on a two-dimensional molecular data storage system that records information in both the sequence and the backbone structure of DNA and performs nontrivial joint data encoding, decoding and processing. Our 2DDNA method efficiently stores high-density images in synthetic DNA and embeds pertinent metadata as nicks in the DNA backbone. To avoid costly worst-case redundancy for correcting sequencing/rewriting errors and to mitigate issues associated with mismatched decoding parameters, we develop machine learning techniques for automatic discoloration detection and image inpainting. The 2DDNA platform is experimentally tested by reconstructing a library of images with undetectable or small visual degradation after readout processing, and by erasing and rewriting copyright metadata encoded in nicks. Our results demonstrate that DNA can serve both as a write-once and rewritable memory for heterogenous data and that data can be erased in a permanent, privacy-preserving manner. Moreover, the storage system can be made robust to degrading channel qualities while avoiding global error-correction redundancy.

## Introduction

DNA-based data storage systems are viable alternatives to classical magnetic, optical, and flash archival recorders^1^. Macromolecular data storage platforms are nonvolatile, readout-compatible, extremely durable and they offer unprecedented data densities unmatched by other modern storage systems^2–10^. Traditional DNA-based data recording architectures store user information in the sequence content of synthetic DNA oligos within large pools that lack an inherent ordering, and user information is retrieved via next-generation or nanopore sequencing^6^. Despite recent progress, several issues continue to hinder the practical implementation of molecular information storage models, including the high cost of synthetic DNA, lack of straightforward rewriting mechanisms, large write-read latencies, and missing oligo errors incurred by solid-phase synthesis.
Image data is typically compressed before being recorded, and even a single mismatch can cause catastrophic error-propagation during decompression and lead to unrecognizable reproductions^6,11,12^. Moreover, the rate of synthesis and sequencing errors may vary an order of magnitude from one platform to another, while PCR reactions and topological data rewriting may cause additional gradual increases in sequencing errors. Therefore, to ensure accurate reconstruction, one needs to account for the worst-case scenario and perform extensive write-read-rewrite experiments to estimate the error rates before adding redundancy^13–15^. Moreover, the estimated error rates have to be accurate enough for efficient error correction due to the mismatched decoding parameter problem^16,17^. The mismatched-decoder problem is an issue mostly overlooked in prior works and it asserts that powerful error-correction schemes such as low-density parity-check (LDPC) codes^18^ require good estimates of the channel error probability to operate properly. This is clearly hard to achieve for traditional DNA-based data storage systems due to the highly stochastic nature of the PCR, sequencing and rewriting process.

Here, we develop and experimentally test a hybrid DNA-based data storage system termed 2DDNA, to address the issue of rewriting and avoid the use of worst-case error-correcting redundancy needed to combat random and missing oligo errors that may accumulate in time and due to content changes. 2DDNA uses two different information dimensions and combines desirable features of both synthetic and nick-based recorders^19^. This is achieved by superimposing metadata (such as ownership information, dates, clinical status descriptions) stored via nicks onto images encoded in the sequence. Sequence content carries large amounts of information, but rewriting is difficult; information stored in nicks^19^ is usually of smaller volume but highly amenable for efficient, permanent and privacy-preserving erasing and rewriting. Importantly, information in both dimensions can be read simultaneously, as locations of nicks are determined using the nick-free strand as reference. Our approach is based on a simple compression scheme for images that operates separately on three different color channels and combines newly developed and existing machine learning (ML) and computer vision (CV) techniques for image reconstruction and enhancement to create high-quality replicas of the original data. For some images with highly granular details, we also propose unequal error protection methods^20^ based on LDPC codes^18^ that only introduce redundancy for sensitive facial features. The 2DDNA paradigm eliminates the need for worst-case coding redundancy and avoids problems with mismatched decoding parameters. It offers the possibility for users to retrieve images of quality dictated by their channel error rates, which may be seen as a form of multiresolution coding. It also offers high information density and simultaneously enables rewriting of data recorded in the backbone via ligation followed by enzymatic nicking, lending itself for use in applications with both synthetic and native DNA substrates for the sequence content^19^.

## Results

### Sequence Dimension Encoding

The encoding framework of 2DDNA is shown in Fig. 1. In the sequence dimension, we perform aggressive quantization and specialized lossless compression that leads to two-fold file size reductions. Compression is known to cause significant losses in image quality when errors are present, so it is common practice to include up to 30% error-correction redundancy^4,7^ which ultimately increases the cost of the storage system. We avoid error-correction redundancy and instead tailor our compression algorithm to accommodate image processing techniques from ML and CV to restore the image to its original quality. The specialized encoding procedure involves two steps, depicted in Fig. 1a. First, RGB channel separation is followed by 3-bit quantization and separate lossless compression of the three color channels. The latter process is performed using the Hilbert space-filling curve^22^ (Supplementary Fig. 1) which preserves local 2D image similarity and smoothness, thereby resulting in linear strings with small differences between adjacent string entries. Moreover, we further employ differential encoding^23^ that involves taking differences of adjacent string values to create new strings with a high probability of small symbols. Differential encoding is followed by Huffman encoding^24^ which exploits the bias towards small symbol values. Together, these operations are performed separately on strings partitioned into eight subsets according to their quantized intensity (brightness) levels. Note that in our ML-based image reconstruction approach, we do not try to optimize the compression scheme: One may also use a basic 3 -bit quantization scheme without lossless compression, at the cost of slightly increased file sizes. Results pertaining to this approach are described in the Supplementary Information (SI), Supplementary Discussion.

**Figure 1.**
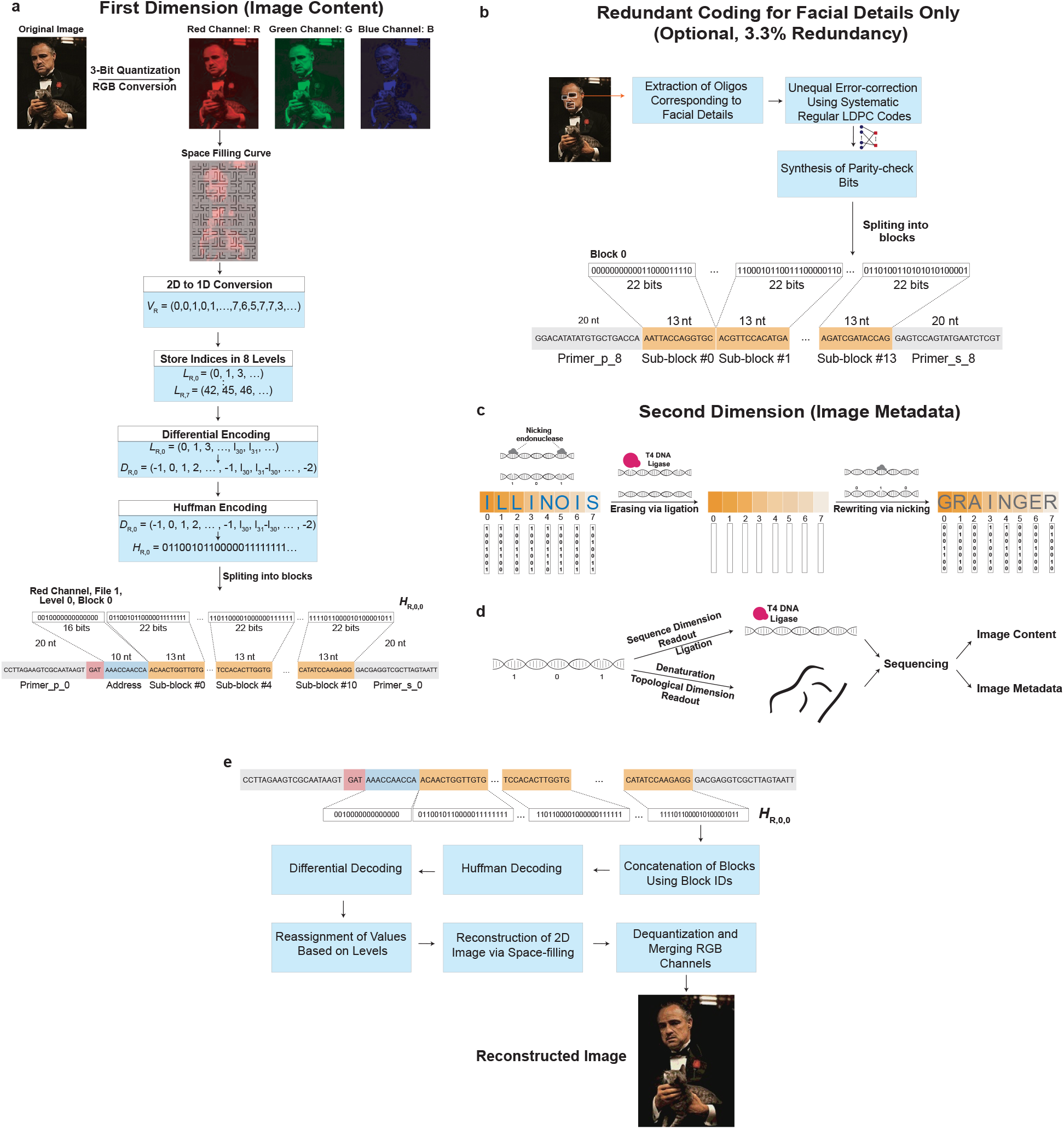
Schematic of the encoding and decoding procedure of our 2DDNA system. **a**) The encoding procedure in the first dimension (sequence content) entails splitting the color image into the Red (R), Green (G) and Blue (B) channel; aggressively quantizing the RGB channels from 256 to 8 intensity levels; performing lossless compression of individual channels through a combination of 2D to 1D conversion of the image data via space-filling curves followed by differential and Huffman encoding. Note that the encoding procedure is separately applied to each intensity level, and the generated binary vector is further augmented by channel information and addresses used to access the oligos. The scheme does not include error-correction redundancy. **b)** For images with granular and highly relevant image features, one can optionally use unequal error-correction coding based on low-density parity-check (LDPC) codes with only 3.3% redundancy compared to the scheme without redundancy. **c)** The encoding procedure in the second dimension (topological content) entails representing letters of the English alphabet in ASCII format and designating one nicking endonuclease to each of the seven bits in the format. Information is encoded using mixtures of endonucleases for which the ASCII bit is equal to 1. Rewriting is performed by sealing the nicks using the T4 DNA ligase and repeating the previously outlined procedure with different data. **d)** 2D data readout through the use of two subpools, one for each storage dimension. **e**) Image decoding is performed by reversing the steps of the encoding process in the first dimension. The image^26^ used in this figure is courtesy of Paramount Pictures. The original data are provided in the Source Data file.

Our encoding involves a second step that translates the binary strings into DNA oligo sequences. Here, DNA oligos of length 196nts are parsed into the following three subsequences (Fig. 1a): (1) a pair of primer sequences, each of length 20nts, used as prefix and suffix, (2) an address sequence of length 10nts, and (3) 11 information-bearing sequences of length 13nts. Primer and address sequences are used for PCR amplification and random access^5^. In addition, a block of three nucleotides is prepended to the address sequence to represent the RGB color information. When converting binary data into DNA sequence content, we use two additional constrained mappings to ensure that the maximum run length of G symbols is limited to three (to avoid G quadruplexes), and that the GC content is in the range of 40 – 60%. Overall, the mapping scheme converts blocks of 16 bits into blocks of 10nts for the address sequences, and blocks of 22 information bits into blocks of 13nts. A detailed description of each step, including the addition of synchronizing markers, is provided in the SI, Supplementary Methods.

### Topological Dimension Encoding

In the topological dimension, we record the metadata in nicks created on the backbone of the synthetic DNA molecules by transforming and generalizing our Punch-Cards system^19^ that was also used for specialized in-memory molecular computing^25^. The main modifications consist in disposing of nicking enzymes that require the additional synthesis of specific guide sequences; native nicking endonucleases are used instead by employing ON-OFF encoding across different intensity pools. Short binary strings are converted into combinations of native nicking endonucleases that determine the composition of nicked/unnicked sites. More precisely, a set of complementary nicking endonucleases is used as the writing tool and selected based on two main criteria: (1) endonucleases must be highly site-specific to prevent non-specific cleavage of the DNA template and hence preserve DNA integrity; and (2) recognition sequences should be selected with sufficiently large Hamming distances between them to prevent undesired cross-nicking (i.e., an enzyme nicking an undesired target site). The mixture composition determines which letter is stored based on the corresponding ASCII code, with the caveat that a ‘1’ is encoded through the presence of the enzyme in the mixture (ON), whereas a ‘0’ is encoded through the absence of the enzyme (OFF). This method enables superimposing information on top of data stored in the DNA sequence content, with no need to change the synthetic platform, as shown in Fig. 1c. Nevertheless, it introduces readout challenges as the nicks break the structure of the strands and may hence lead to assembly ambiguities. We address this problem via an algorithmic solution that involves searching for potential prefix-suffix substrings in the nicked pool.

### DNA Synthesis and Sequencing

To demonstrate a proof-of-concept, we experimentally tested the storage platform on eight Marlon Brando movie stills, shown in Fig. 2a. The original files were of total size 8,654,400bits, but after the two-step encoding procedure (Fig. 2b), they reduced to 2,317,896nts. The corresponding 11,826 DNA oligos were synthesized by Integrated DNA Technologies (IDT). One pool was reserved for each of the eight levels. The oPools were sequenced on an Illumina MiSeq device following standard protocols described in the Methods. Individual sequence reads may contain errors, so we first construct a consensus sequence by aligning reads with error-free addresses, following the approach described in our prior work^6^. This process led to 11,726 perfectly recovered sequences and 22 sequences that contain errors but do not significantly compromise the image quality; 78 oligos were either highly corrupted or completely missing from the pool.

**Figure 2.**
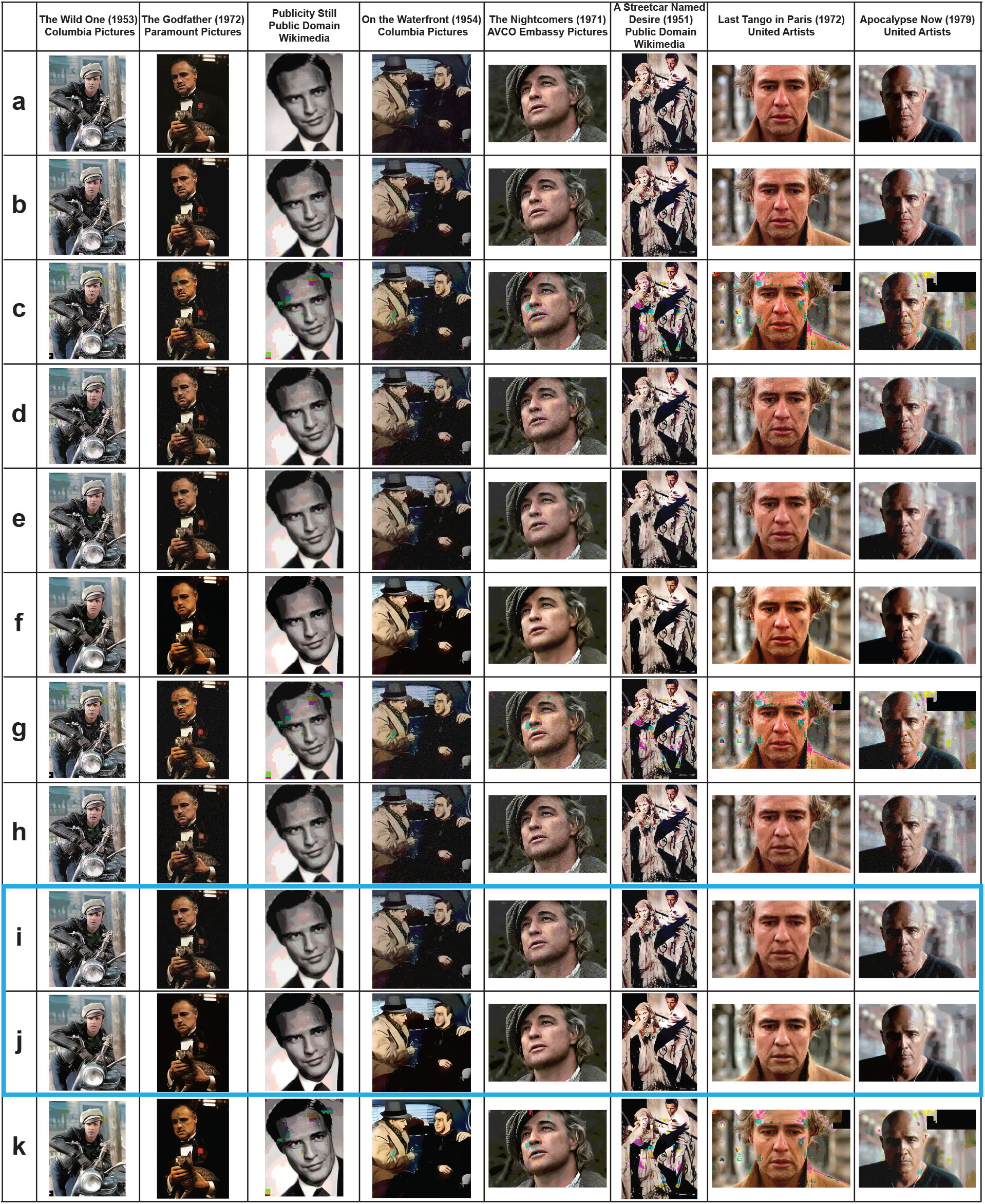
Write-Read results for encoding information content in the sequence dimension. **a**) Original images with 256 RGB intensity levels, encoded by 8 bits each. **b**) Quantized images with 8 RGB intensity levels, encoded by 3 bits each. **c**) Images generated directly from the information encoded in DNA oligos without error-correction redundancy. **d**) Images reconstructed after applying a combination of discoloration detection and image inpainting on the results in **c**. **e**) Images refined via smoothing of the results depicted in **d**. **f**) Image enhancement results for images shown in **e. g**) Images reconstructed using unequal error-correcting coding for facial features. **h**) Images reconstructed after applying a combination of discoloration detection and image inpainting on the results in **g**. **i**) Images refined via smoothing of the results depicted in **h**. **j**) Image enhancement results for images shown in **i**. **k**) Image enhancement results for images shown in **c**. In summary, the best quality results - obtained using our image processing techniques - are given in **i** and **j** (boxed). The images in this figure are courtesy of: Paramount Pictures, Sony Pictures, MGM Studios, StudioCanal, American Zoetrope (© 1979 Zoetrope Corp. All Rights Reserved.), the Marlon Brando and Rod Steiger estates. The black and white public domain still of “A Streetcar Named Desire” was colorized using the software Hotpot.ai.

### Sequence Dimension Decoding and Post-Processing

The images generated from this procedure are depicted in Fig. 2c. Upon close inspection, it is apparent that the encoded images suffer from visible degradation, and in particular, large blocks of discolorations. These artifacts can be removed by applying a carefully designed combination of ML and CV image processing techniques (Fig. 4), tailor-made to operate on images compressed according to our method.

**Figure 3.**
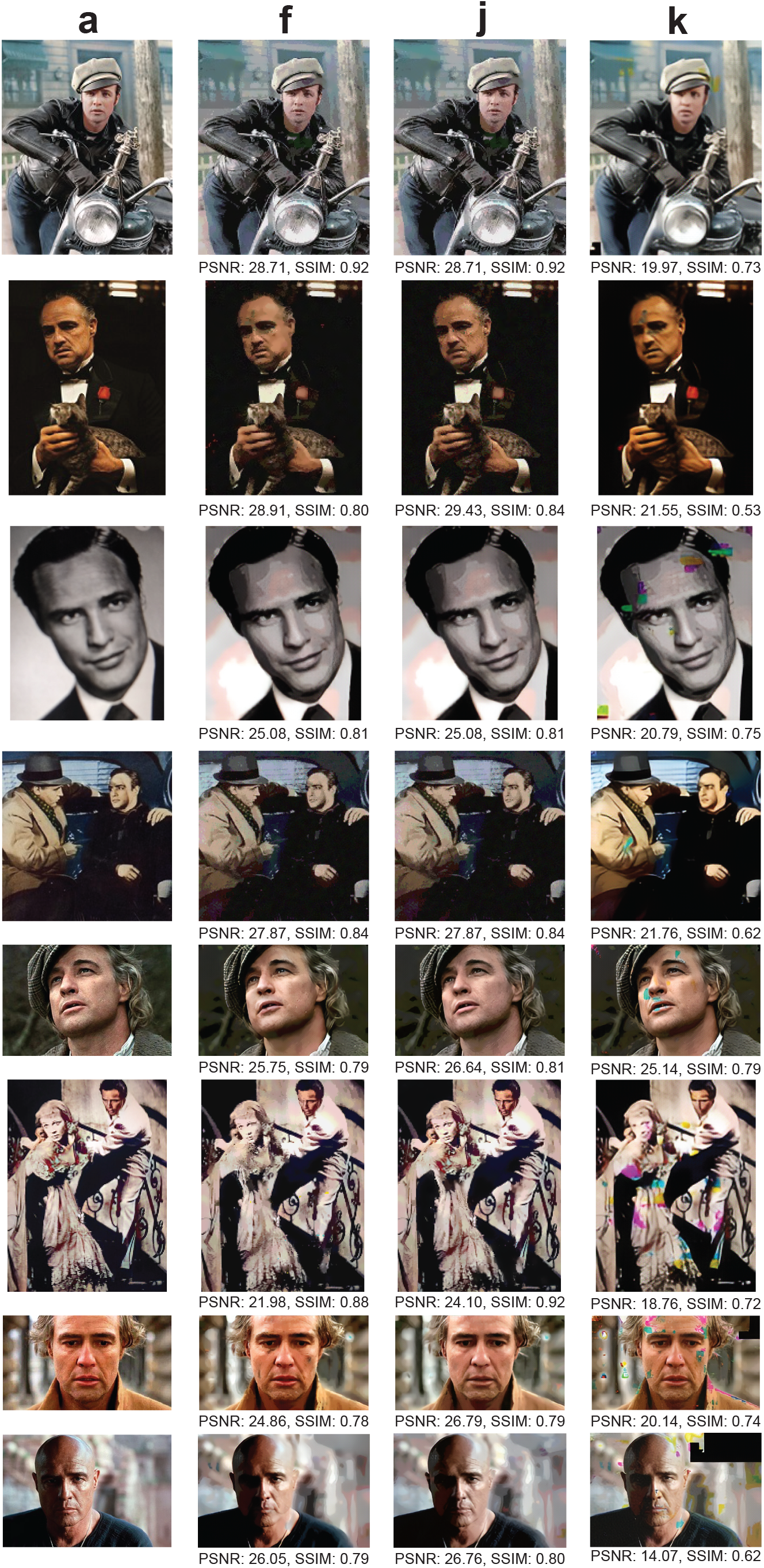
Comparison of our automatic discoloration detection and inpainting approach with the state-of-the-art image enhancement technique^33^. The results shown include quantitative performance metrics computed with respect to **a**. The column labels refer to the corresponding rows in Fig. 2. Column **a**): The original, uncompressed images. Column **f**): The images reconstructed using our method, without unequal protection redundancy for facial features. Column **j**): The images reconstructed using our method, with roughly 3.3% redundancy for facial features. Column **k**): Results obtained after image enhancement, applied directly to the decoded DNA oligo images with errors. The pictures in this figure are courtesy of: Paramount Pictures, Sony Pictures, MGM Studios, StudioCanal, American Zoetrope (© 1979 Zoetrope Corp. All Rights Reserved.), the Marlon Brando and Rod Steiger estates. The black and white public domain still of “A Streetcar Named Desire” was colorized using the software Hotpot.ai. The original data are provided in the Source Data file.

**Figure 4.**
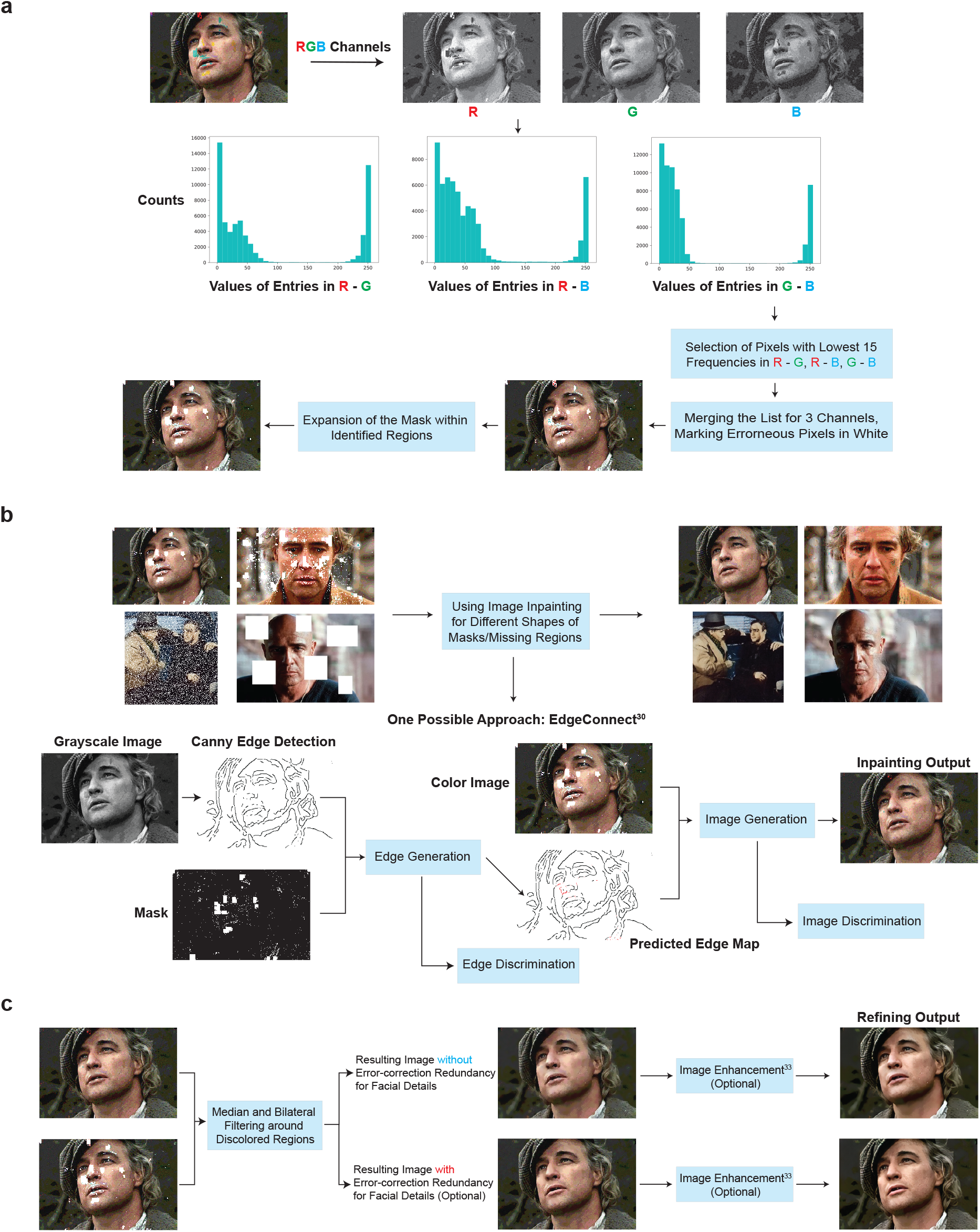
Diagram of the ML postprocessing techniques used to reconstruct images encoded in oPools. **a)** Automatic discoloration detection based on the natural redundancy in the three RGB color channels. The histograms reflect the frequency counts of the pairwise differences in channel intensity levels which are used to assess which color channel may contain errors (SI, Supplementary Methods). **b**) Pixel masking and inpainting via deep-learning architectures. **c)** The smoothing and image enhancement procedures. The pictures in this figure are courtesy of: Paramount Pictures, Sony Pictures, MGM Studios, StudioCanal, American Zoetrope (© 1979 Zoetrope Corp. All Rights Reserved.), the Marlon Brando and Rod Steiger estates. The original data are provided in the Source Data file.

Note that our compression scheme mitigates the effects of catastrophic error-propagation which may be otherwise present when using a JPEG compressor (Supplementary Fig. 6). As JPEG formats are highly sensitive to errors, they result in poor-quality reconstructions if one does not use a coding overhead that guarantees exact reconstruction. Furthermore, alternative methods based on joint source-channel coding^41^–^42^ still require introducing error-control redundancy which we are aiming to dispose of in our learning-based approach. To demonstrate this point, we performed extensive simulations with six combinations of JPEG image compression qualities and matching error-control coding schemes. For JPEG-compressed files with different quality parameters (as defined in the Python Pillow Package for all image formats, JPEG included, on a scale from 1 (worst) to 95 (best)), we added LDPC redundancy to the compressed data for error-correction to ensure that the resulting number of oligos (file size) is as close as possible to that used in our experiment. The base substitution error is set to 0.8%, while the missing oligo error is set to 0.7%, matching the numbers obtained experimentally, leading to an overall bit error of 1.9%. We decoded the binary information from the erroneous DNA oligos using LDPC codes, followed by JPEG reconstruction. Part of the results are shown in Fig. 5 and full set of results are shown in Supplementary Fig. 7. Note that since JPEG has very specific formatting rules, missing or erroneous critical identifiers in JPEG files leads to system errors, such as OSError in Python. Other compression methods, such as those based on Generative Adversarial Nets (GANs) are discussed in the SI, Supplementary Discussion and Supplementary Fig. 10.

**Figure 5.**
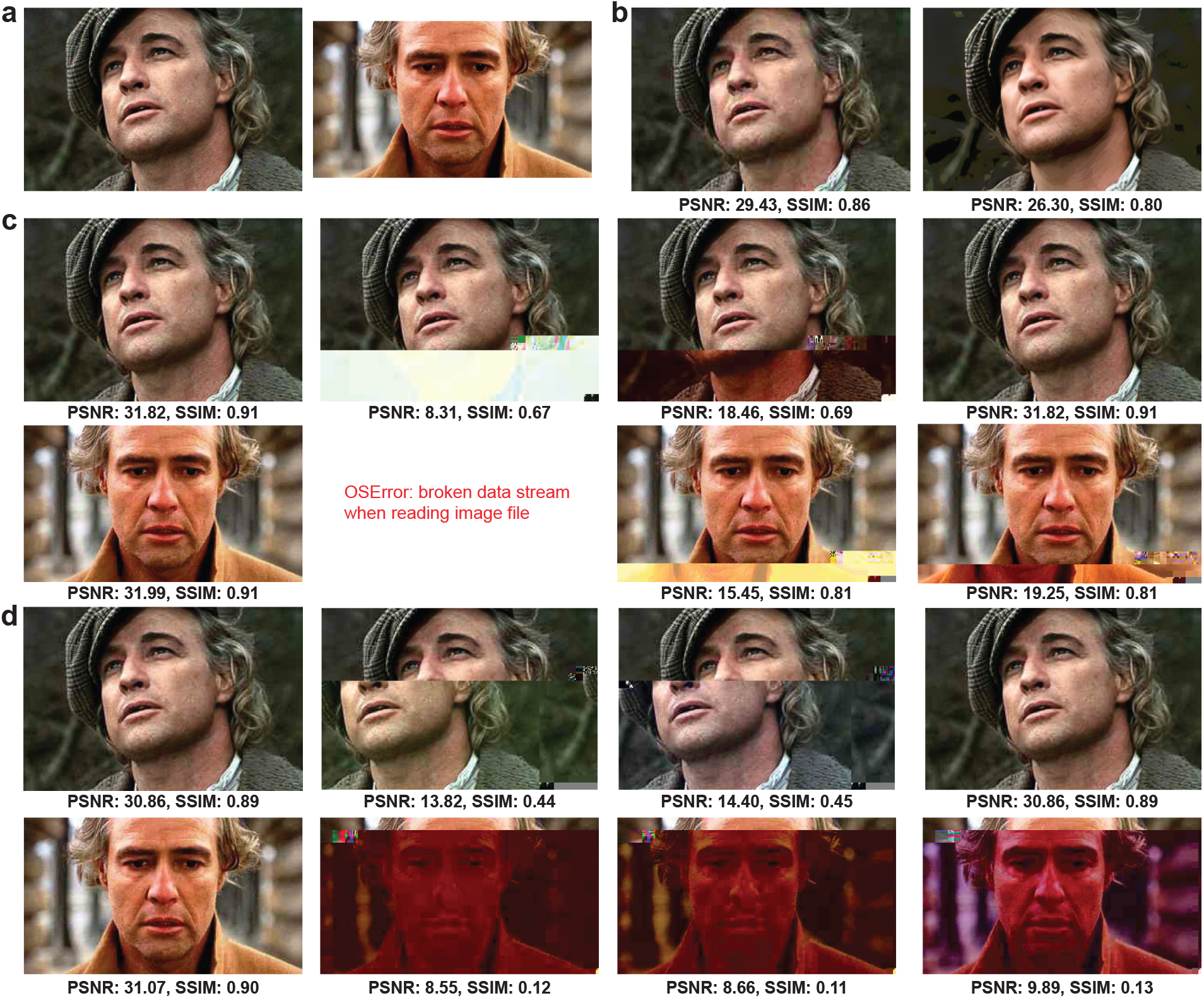
a) The original, uncompressed images. **b**) An example illustrating why single-value metrics for assessing image quality are inadequate. Left: An image compressed by JPEG with quality parameter 20. Right: The same image, after simple 3-bit quantization and image enhancement^33^. The image on the right is visually superior yet has consistently worse numerical quality metrics compared to the image on the left. **c**) Images are compressed with JPEG quality parameter 40 and encoded with an LDPC code of rate R=0.125. The two images in the first column: No errors are added, so the decoding procedure is successful. Images in the remaining three columns represent pairs of decoded images with LDPC channel error rate parameters (probabilities) 0.5%, 1%, and 5%, respectively; in these cases, the base substitution error equals 0.8%, the missing oligo error rate equals 0.7%, resulting in an overall bit error rate of 1.9%. The channel parameter for LDPC decoding is assumed not to be known beforehand. **d**) Images compressed with JPEG quality parameter set to 30 and encoded with an LDPC code of rate R=0.108. The two images in the first column: No errors are added, so the decoding procedure is successful. Images in the remaining three columns presenting pairs of decoded images with LDPC channel error rate parameters (probabilities) 0.5%, 1%, and 5%, respectively; in these cases, the base substitution error equals 0.8%, the missing oligo error rate equals 0.7%, resulting in an overall bit error rate of 1.9%. The pictures in this figure are courtesy of: Paramount Pictures, Sony Pictures, MGM Studios, StudioCanal, the Marlon Brando and Rod Steiger estates. The original data are provided in the Source Data file.

To correct for image discolorations, we implement a three-step post-processing procedure that has no matching counterpart in the digital domain and heavily relies of using the color channels as a natural source of redundancy. The first step includes detecting the locations with discolorations and masking them out, as shown in Fig. 4a and Supplementary Fig. 3. To pinpoint the discolored regions without direct visual inspection (i.e., in an automated fashion), as already pointed out, we leverage the separate information content in the three distinct RGB color channels. Due to the random nature of errors, it is highly unlikely to have correlated errors in multiple channels for the same pixel. Hence, the three-color decomposition acts as a 3-repetition cod, GatedConvolutione because at least two of the three color channels are likely to be unperturbed. A detailed explanation of the technique can be found in the SI, Supplementary Methods, which is adapted from our preliminary approach^27^. The second step involves using an existing deep learning technique known as image inpainting^28–30^ to replace the masked pixels with values close to the original. Neural networks are well-suited for inpainting because they can be trained on massive datasets. For our system, we use the state-of-the-art GatedConvolution^28^ and EdgeConnect^30^ methods. The basic architecture of EdgeConnect is shown in Fig. 4b and Supplementary Fig. 4, and the results after applying discoloration detection and image inpainting are shown in Fig. 2d. Finally, the third step involves smoothing the image to reduce blocking effects caused by quantization and blending mismatched inpainted pixels, as shown in Fig. 4c and Supplementary Fig. 5. Here, we use bilateral^31^ and adaptive median smoothing^32^ on the coarsely inpainted images, and we include additional image enhancement features^33^ to further improve image quality. The image post-processing procedure relies on storing R, G and B color channels in different oligos and using the channels as “proxies” for repetition codes. This ensures that it is highly unlikely to have correlated errors in multiple channels for the same pixel and that discolorations can be detected through majority rules. As a result, our scheme can be used with any other type of recorder that splits images into R, G, B subimages and stores them separately.

The results of image smoothing are depicted in Fig. 2e, and the enhanced images are shown in Fig. 2f As shown in Figs. 2e, f and Supplementary Fig. 2, some facial details in highly granular images remain blurred even after applying the learning methods. To address these issues, we further propose the use of unequal error-protection for such images, which implies adding highly limited redundancy only to oligos bearing facial features (e.g., eyes, lips), as shown in Fig. 1b and explained in the SI, Supplementary Methods. Redundancy is added through a regular systematic LDPC codes of rate 0.75, resulting in 391 additional oligos and an overall overhead of 3.3%. Images generated from this redundant pool are shown in Fig. 2g, whereas Figs. 2h, i, j parallel the results of Figs. 2d, e, f for the case of no unequal error-correction redundancy. Note that there exist no other approaches to performing the same task in the signal processing and computer vision community. Applying state-of-the-art image enhancement method^33^ directly on images generated from error-bearing DNA oligos without error-correction results in poor quality reconstructions because classical image enhancement methods cannot automatically correct discolorations (Fig. 2k and Fig. 3k). Both quantitative metrics (Peak Signal-to-Noise Ratio (PSNR) and Structural Similarity (SSIM)) as well as visual inspection of the recovered images show that our method offers significantly better performance than direct image recovery and enhancement of the corrupted DNA-encoded images. Processed images with corresponding quality values are plotted in Figs. 2a, f, j, k and Fig. 3.

For LDPC codes, it is crucial to have good estimates of the channel error probability: LDPC belief propagation decoding performs well in practice but is highly sensitive to incorrect initial loglikelihood ratios, which are functions of the channel error rate^16,17^. Therefore, when using mismatched channel parameters, LDPC decoders can fail to correct all errors, which in turn can lead to corrupted JPEG decoding, as seen in Fig. 5. It is worth pointing out that correlations amongst errors may cause some oligos to be disproportionally affected and others to have barely any errors. To further mitigate this issue, oligo-level redundancy was used^4^ before, but here it is replaced by a concatenation of an interleaver and LDPC codes, as interleaving renders errors uncorrelated and helps with missing oligo content reconstruction. We present additional results related to LDPC coding with interleaving in the SI, Supplementary Discussion and Supplementary Fig. 8).

### Topological Dimension Recording and Post-Processing

As a proof of concept for storage in the topological dimension, we superimposed information on the same Marlon Brando images (Fig. 7). In the writing experiment, we recorded the word “ILLINOIS,” comprising 56 bits in ASCII code, across eight different intensity-level DNA pools. We selected seven nicking endonucleases, each representing one bit of the 7-bit ASCII code. These enzymes have recognition sites that exist in at least one oligo of each of the eight pools, and the sites are used as recording positions. In the ASCII code, ‘1’ translates into inclusion, whereas ‘0’ translates into exclusion of the corresponding enzyme. Upon nicking, the pools are sequenced using the procedure described in Fig. 1d. In this way, the nicked oligos were denatured, resulting in ssDNA fragments of various lengths dictated by the position of the nicks. The fragments were subsequently converted into a dsDNA library and sequenced via Illumina MiSeq. To verify the existence of short-length fragments capped at both ends by enzyme recognition sites, we developed a detection algorithm with a flowchart depicted in Fig. 6. The gist of the algorithm is to detect if a nick was created or not based on a search for two fragments corresponding to the prefix and suffix of the sequences recognized by the enzyme. Note that our algorithm counts the number of appearances of all possible (potential) nicking events for the sets of enzymes used. The decision regarding which enzymes are included in a certain pool is based on the counts of each prefix-suffix pair. To rewrite the data, we performed the process outlined in Fig. 1c, which involves treatment of the nicked DNA with the T4 DNA ligase. This erasure method completely removed the recorded metadata. Note that the ligase was perfectly effective in so far that each original oligo was accounted for in the sequenced pool. We then rewrote the word “GRAINGER” using the same topological nicking process with error-free reconstruction.

**Figure 6.**
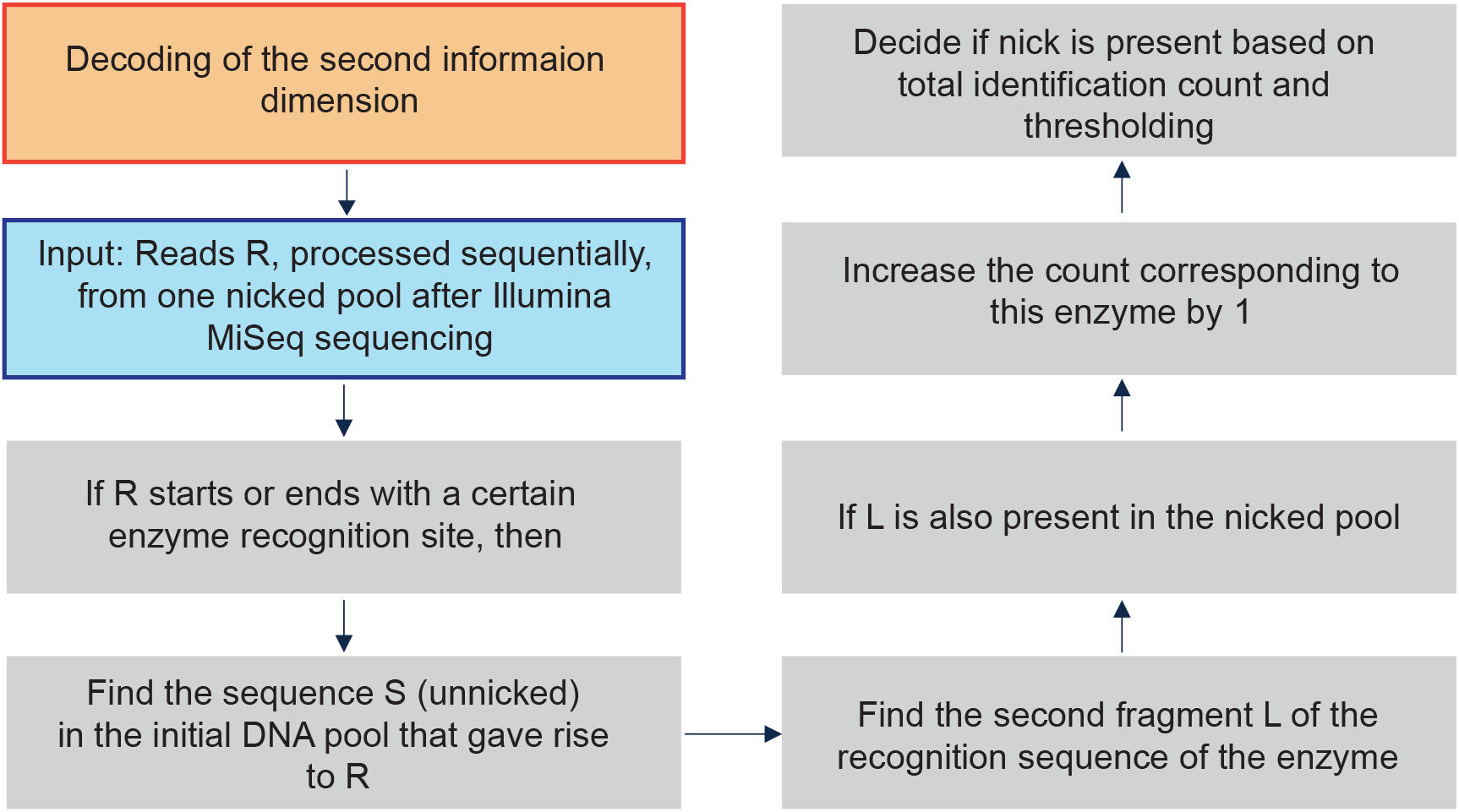
A prefix-suffix pattern search method for decoding the information in the topological dimension. Each fragment obtained from the nicked pool is searched for the presence of a prefix-suffix substring pair that can indicate that a specific enzyme was included in the combinatorial mixture used for the given intensity pool. Since undesired nicking reactions may occur, some counts corresponding to recognition sites of enzymes that were not actually present in the pool may be nonzero; in this case, we make the decision based on how large the counts are relative to others (i.e., we use thresholding with a threshold determined based on the largest count values for the pool).

**Figure 7.**
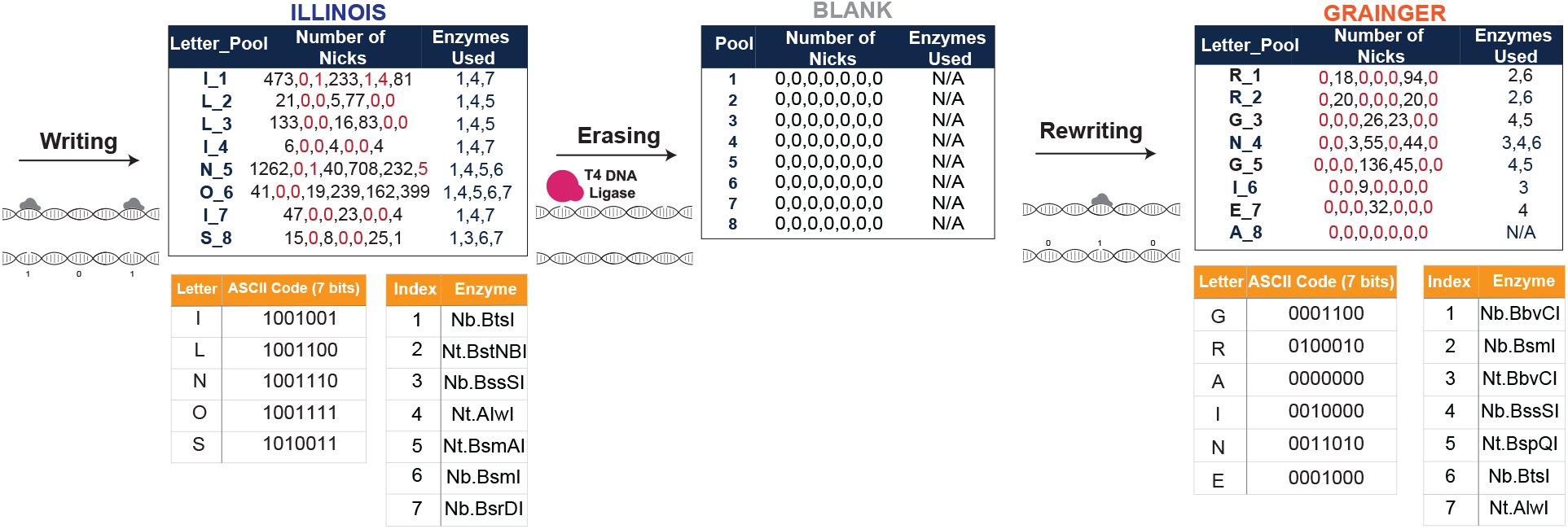
Schematic of metadata encoding and identification using DNA nicking. The numbers in the middle column of the leftmost and rightmost tables represent the number of oligos in the sequenced pool capped by the recognition sequences of the nicking enzymes. The numbers listed in red correspond to the labels of nicking enzymes not used in the encoding of the letter to the left. As may be seen from both tables, the largest red numerical value is significantly smaller than the smallest black value for all encodings (e.g., 4<<81, 5<<40 in the leftmost table) and the second round of writing resulted in no spurious nicks whatsoever. The quality of the results in the rewriting experiment may be attributed to a more suitable choice of nicking enzymes determined upon inspection of the results of the first round. Hence it is recommendable to use the second collection of enzymes for recording purposes. Also note that we shuffled the symbol encodings for the rewriting experiment in order to test more combinations of nicking enzymes. In the erasure step, the T4 DNA ligase was used in a single step reaction to seal all the nicks. No nicks were found after the ligation reaction, showing that the ligase perfectly erased the data (middle table). Note that when recording “ILLINOIS” and “GRAINGER” only six and five enzymes were effectively used for the ASCII code, respectively, due to the choice of the letters in the words.

As outlined above, decoding the information stored in the two dimensions requires nontrivial approaches, involving new pattern search algorithms. To hence read the content stored in both dimensions, two separate subpools are retrieved for each level. The sequence content is reconstructed by first sealing the nicks in one of the two subpools via ligation, as done during rewriting, followed by sequencing. Alternatively, to avoid ligation for the sequence content readouts, one may choose to only record the topological information on a subpool of oligos. This resolves the problem of sorting the nicked oligo fragments. The content in the nicks is retrieved using the second subpool. After sequencing, the reads are aligned to the now known full-length reads obtained from the first subpool in which the nicks were sealed. The results of the alignment are used in the algorithmic procedure to determine which enzymes were used for nicking and consequently, for reconstruction of the ownership metadata (Fig. 7).

## Discussion

Existing technologies for DNA synthesis, editing, and sequencing allow for writing and reading diverse information in multiple dimensions or molecular features. Our 2DDNA platform exploits these tools to enable recording data in two DNA dimensions, including sequence context and backbone structure, thereby opening the door for multidimensional macromolecular storage systems that can use multiple molecular properties (including molecular concentration). Our results show that the 2DDNA system takes advantage of our automatic discoloration detection approach and powerful state-of-the-art deep learning methods for image inpainting and enhancement to substantially improve the quality of the stored images without error-control redundancy. This represents a fundamental advancement in molecular storage which departs from prior techniques in the field and reduces the cost of data storage by greatly minimizing or eliminating the need for synthesizing redundant oligos. The tailor-made learning methods also overcome reliability issues that cannot be addressed by off-the-shelf JPEG compression and joint source-channel coding methods.

Our storage system also offers a simple means for permanently erasing metadata information. The ligation-based approach differs substantially from the existing rewriting methods^5,43^. In the first setting, overlap-extension PCR is used to rewrite blocks of texts corresponding to words. This is a tedious, multi-step approach and much more complex to perform than ligation. In the second approach, one requires additional DNA synthesis and multiple hybridization and strand displacement steps to rewrite the content. Note that in our system metadata is automatically sequenced during the sequencing of the actual image – no separate sequencing for the nick-based information is needed. This is the case since we can always use the strand that is free of nicks as reference for sequence alignment to determine the locations (positions) of the nicks.

For selective amplification and PCR-based random access, the oligos we used to store image content contain carefully designed primers. The primers satisfy Hamming distance, sequence correlation, sequence balance and so-called primer-dimer constraints^44^. Note that once nicks are added to the sugar-phosphate backbone, one cannot run PCR reactions on the oligos directly. To randomly access an image, a certain amount of DNA from the oPools has to be isolated, sealed using the T4 ligase and then amplified via PCR. Consequently, metadata is removed from the selected subpool to enable random access to the image itself, but it remains intact in the global pool of oligos. In order to avoid first sealing the nicks and then running the PCR, one can also use other methods for random access, involving magnetic beads with attached primers corresponding to the address sequences of the image of interest^45^.

Our 2DDNA platform was tested on eight images of total size 1.082MB. The only oligo content that does not correspond to actual raw image information includes primers, pixel/color/image identifiers, constrained redundancy for balancing the GC content and removing long runs of Gs as needed for synthesis. The average sequencing coverage used is 112x, which is small compared to the 3000x coverage reported in^2^ and the 370x coverage from^4^. It is higher than the coverage of 5x reported in^15^ but in that case, error-control coding redundancy is used. We did not try to optimize the sequencing coverage - our coverage values are dictated by the sequencing protocol used and are not needed for high-quality reconstruction. Supplementary Fig. 9 in the SI shows that low-coverage and hence high error-rates can be accommodated within our system, even when the error rate is as high as 7%.

The information density of our platform equals the number of bits stored divided by the number of nucleotides used for encoding. Since quantization is used during the encoding procedure, there are two ways to compute this density: If calculated with respect to the number of bits in the raw image files, the information density equals 3.73bits/nt. Clearly, this exceeds the maximum 2bits per nucleotide density dictated by the 4-alphabet size, but may be seen as a consequence of the fact that we get a distorted image back, which allows for an increase from 2 to 3.73bits/nt. If the information density is calculated with respect to the number of bits of the quantized image files, the information density equals 1.40bits/nt. The reason why this value is smaller than 1.57bits/bp reported in^5^ and 1.72bits/bp reported in^6^, is that in the latter two works gBlocks of length 1000bps were used, while in this work we used oPools of length 196nts. To allow for random access, one has to include primers and address sequences which amount to 53nts per oligo, i.e., per 196 nucleotides - an overhead of 27%. This is to be compared to roughly 50bps per 1000bps^5,6^, resulting in a significantly smaller overhead of 5%. When converted into bytes/gram, the two reported densities theoretically equal 0.91 zettabytes/gram and 0.34 zettabytes/gram.

In conclusion, 2DDNA provides the foundations for storage of heterogeneous datasets with rewriting capabilities and at the same time empowers the use of DNA media for nontraditional applications such as parallel in-memory computing.

## Acknowledgements

The work was funded by the DARPA Molecular Informatics Program, the NSF+SRC SemiSynBio program under agreement number 1807526 and NSF grants 1618366 and 2008125. The stills used in this work are courtesy of: Paramount Pictures, Sony Pictures, MGM Studios, StudioCanal, American Zoetrope (© 1979 Zoetrope Corp. All Rights Reserved.), the Marlon Brando and Rod Steiger estates. Two additional images encoded in DNA are available in our Figshare repository, while the results on Marlon Brando publicity still and the public domain still of “A Streetcar Named Desire” were simulated using a noise model generated from actual DNA-encoded images. The public domain still was colorized using the software Hotpot.ai.

The authors are also grateful to Laura Sevier and Alonzo Wickers for their invaluable help in securing the copyrights for the movie stills.

## Author Contributions

C.P., SMH.T.Y. and O.M. conceived the ML image compression, inpainting and unequal-error protection scheme and performed detailed data analysis. SK.T., C.M.S., SMH.T.Y. and O.M. designed the nicking experiments while SK.T. performed the experiments. A.G.H. performed the sequencing experiments and helped with the data analysis. All authors contributed towards the system development and participated in the writing of the manuscript.

## Data availability

The sequencing and image data generated in this study have been deposited in Figshare repository at: https://doi.org/10.6084/m9.figshare.17162546.v1. Source data, used to create the figures, accompany the online version of this article.

## Code availability

All scripts are available in Zenodo repository at: https://doi.org/10.5281/zenodo.5774385.

## Methods

### oPool PCR Amplification and Sequencing

The list of primers used in our experiments is shown in Supplementary Table 1. The oPools and corresponding primers were ordered from Integrated DNA Technologies (IDT): https://www.idtdna.com/pages/products/custom-dna-rna/dna-oligos/custom-dna-oligos/opools-oligo-pools.

All oPools were diluted to 5ng/ul. The primers were diluted to 10uM. Each oPool was amplified in separate reactions using forward and reverse primers for each of the 8 levels. Reactions were set up with 5ng of oPool, 1ul of each forward and reverse primer diluted to 10uM, 22ul of water and 25ul of Kapa HiFi DNA Polymerase (Roche, CA) with the following PCR cycling conditions: denaturation at 98°C for 45s, 8 cycles of 98°C for 15s, annealing at 51°C for 30s and extension at 72°C for 30s, followed by a final extension at 72°C for 1min and hold to 4°C.

After PCR, the individual reactions were cleaned up with 50ul of AMPure beads (Agilent, CA) and eluted in 20ul of 10mM Tris. The PCR products were quantitated with the Qubit 3.0 fluorometer and run on a Fragment Analyzer (Agilent, CA) to determine the presence of a band of the correct size and the absence of free primers or primer-dimers. The PCR products from each level were pooled in equimolar concentration and the pool was converted into a sequence-ready library with the Kapa Hyper Library Construction kit (Roche, CA) with no PCR amplification. The final library was quantitated with Qubit and evaluated in a Fragment analyzer and further quantitated by qPCR. The library was loaded on a MiSeq (Illumina, CA) and sequenced for 250 cycles from each end of the library fragments with a Nano V2 500 cycles kit (Illumina). The raw fastq files were generated and demultiplexed with the bcl2fastq v2.20 Conversion Software (Illumina).

### ssDNA Nicking Products Preparation for MiSeq Sequencing

All nicked products were purified using the Qiaquick PCR purification kit (QIAGEN) and eluted in ddH_2_O. They were then denatured at 98°C for 5min and immediately cooled down to 4°C. The ssDNA samples were first quantified via the Qubit 3.0 fluorometer. Next, the Accel-NGS^®^ 1S plus DNA library kit (Swift Biosciences) was used for library preparation following the manufacturer’s recommended protocol. Prepared libraries were quantitated using Qubit and then run on a DNA Fragment Analyzer (Agilent, CA) to determine fragment sizes, pooled in equimolar concentration. The pool was further quantitated by qPCR. All steps were performed for each sample separately and no nicked DNA samples were mixed. The pooled libraries were loaded on an MiSeq device and sequenced for 250 cycles from each end of the library fragments with a Nano V2 500 cycles kit (Illumina). The raw fastq files were generated and demultiplexed with the bcl2fastq v2.20 Conversion Software (Illumina).

### Nicking Experiments

The list of enzymes used in our experiments is shown in Supplementary Table 2. 1μg of each amplified library pool was mixed with the appropriate nicking enzymes, determined based the content being encoded and was incubated in proper buffer conditions and temperature for 1h based on the manufacturer’s protocols available at: https://www.neb.com/products/restriction-endonucleases/hf-nicking-master-mix-time-saver-other/nicking-endonucleases/nicking-endonucleases. SnapGene Viewer 5.1.7 was used to visualize DNA sequences and detect nicking sites.

### Machine Learning and Computer Vision Methods

A detailed description of our compression algorithms and the supporting automatic discoloration detection, inpainting, smoothing and enhancement methods is relegated to the SI, Supplementary Methods, due to the technical nature of the methodology used.

