## Supplementary Information for "Rewritable Two-Dimensional DNA-Based Data Storage with Machine Learning Reconstruction"

### Table of Contents

**Supplementary Methods**

- Encoding Procedure for Sequence Dimension (3-7)
- Decoding Procedure for Sequence Dimension (8)
- Post-Processing for Sequence Dimension (9-12)

**Supplementary Discussion**

- Additional Results on ML-Based Reconstruction versus JPEG with LDPC Coding Redundancy (13-18)
- Additional Results on JPEG versus GAN-Based Compression (19)
- Error Rate and Information Density (20)

**Supplementary Tables**

- List of Used Primers (21)
- List of Used Enzymes (22)

**Supplementary References** (23)

### Supplementary Methods

#### 1. Encoding Procedure for Sequence Dimension

Our two-step encoding procedure first translates an image file into 24 binary strings (if 3-bit quantization is used), and then converts the binary strings into DNA oligos for storage and amplification. A detailed description of each step used in the process is provided below.

##### 1.1 Quantization and Conversion into 2D Color Arrays

The first step in the procedure is RGB channel separation and quantization. To begin with, we split the color images into three color channels, R (red) G (green) B (blue), and then perform 3-bit quantization of the values in each channel. More precisely, the image $I$ is represented by a three-dimension tensor of size $M \times N \times 3$, which is later split into three matrices $R, G, B$, each of size $M \times N$. Next, we perform quantization of each color matrix, leading to the intensity values mapped from $[0,1,\dots,255]$ to $[0,1,\dots,7]$. Specifically, we use the following quantization rule for all three channels:

$$Q[m,n] = \left\lfloor \frac{X[m,n] \times 8}{256} \right\rfloor, m = 1,2,\dots,M; n = 1,2,\dots,N,$$

where $Q$ is the quantized matrix and $X \in \{R, G, B\}$.

##### 1.2 Conversion of 2D Arrays into 1D Vectors

There exist several methods for converting a matrix into a vector so as to nearly-optimally preserve 2D image distances in the 1D domain, such as the Hilbert space-filling curve[1]. The Hilbert space-filling curve, shown in Supplementary Fig. 1, provides a good means to capture 2D locality and is the method of choice in our conversion process. Note that the Hilbert curve is commonly used on square matrices, so we adapt the transversal implementation by recursion to account for matrices with arbitrary dimensions. After the mapping, the matrices $R, G, B$ are converted into vectors $V_R, V_G, V_B$, respectively.

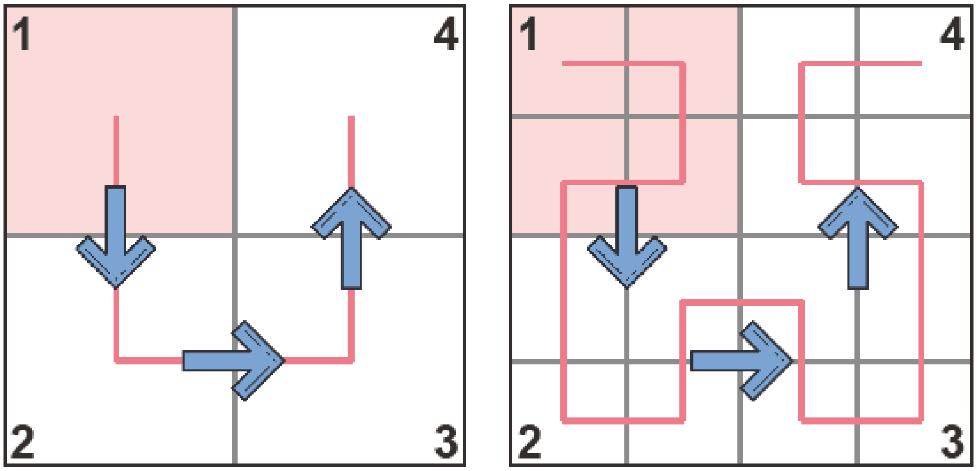


**Supplementary Figure 1.** Hilbert space-filling curves for 2 × 2 and 4 × 4 squares.

#### 1.3 Partitioning of Vectors According to Levels

Upon quantization, the values in $V_R, V_G, V_B$ lie in $[0,1,\ldots,7]$. We next decompose each vector into subvectors of possibly different lengths according to the intensity value. Specifically, $V_R$ is decomposed into $L_{R,0}, L_{R,1}, \ldots, L_{R,7}$, where the vector $L_{R,j}$ contains the indices of the elements in $V_R$ whose value equals $j$. The same procedure is also performed for the vectors $V_G, V_B$. An example decomposition may read as:

$$V_R = 0,0,0,1,7,6,7,7,\ldots \rightarrow L_{R,0} = 0,1,2,\ldots$$

Note that the elements in $V_{X,j}$ are assigned to $L_{X,j}$ in order, where $X \in \{R, G, B\}, j \in \{0,1,\ldots,7\}$. Hence each vector $L_{X,j}$ will only contain increasing values, a fact that we exploit in our later design. Given the Hilbert scan, one may expect the differences between adjacent entries in each of the vectors to be small with high probability. Therefore, splitting a vector into individual levels enables subsequent differential encoding. After the RGB decomposition, quantization and level partition, each image is represented by 24 vectors.

#### 1.4 Lossless Compression

##### 1.4.1 Differential Encoding

Differential encoding[2] converts a vector into another vector containing the initial value of the original and the differences between consecutive values, summarized in vectors denoted by $D_{X,j}$. In order to prevent catastrophic error propagation caused by few errors in encoded vector, we set around 3% of the values in each of these $D_{X,j}$ to their original undifferentiated values and prepend to them the value $-1$. We also append an additional value $-2$ to $D_{X,j}$ to indicate the end of the vector. Under such construction, a typical pair of $L_{X,j}$ and $D_{X,j}$ may read as:

$$L_{X,j} = l_1, l_2, \dots, l_{31}, l_{32}, \dots \rightarrow D_{X,j} = -1, l_1, l_2 - l_1, \dots, -1, l_{31}, l_{32} - l_{31}, \dots, -2$$

Note that as $L_{X,j}$ only contains increasing values, the additional synchronizing markers $-1$s and $-2$s will not be confused with other information-bearing values in $D_{X,j}$. In the decoding procedure $-1$s serve as markers to indicate the ground truth positions and $-2$s represent the end of the vector. One can choose whether to perform run length coding or not after differential encoding based on the statistics of the obtained differential values.

##### 1.4.2 Huffman Encoding

Huffman encoding[3] is performed after differential encoding, where all values in $D_{X,j}$ are used to construct the Huffman code dictionary. This results in a collection of binary strings $B_{X,j}$.

#### 1.5 Conversion of Binary Strings into DNA Oligos

Because the maximum length of high-quality synthetic DNA is constrained, we split the binary strings $B_{X,j}$ further into different substrings to be able for conversion. If needed, the last substring will be padded with dummy values to ensure uniform lengths. Again, the marker $-2$ is used to indicate the end of the vector, so the dummy values can be easily ruled out during decoding procedure. These binary substrings are then converted into oligo sequences over the alphabet $A, T, C, G$ of length 196nts. Each oligo can be parsed into the following three subsequences. The first part is a pair of primer sequences, each of length 20nts, used as prefix and suffix. They allow for PCR amplification and random access. The second part is the address, which records information about the image and its position of corresponding substring in the original binary string. Note that a block of three nucleotides (R: GAT, G: TCG, B: ATC) is prepended to the address sequence to represent the RGB color information. The total length of address sequence is 13nts. The last part is a collection of 11 information-bearing sequences of length 13nts. As mentioned in the main text, the conversion from binary strings to DNA oligos is accomplished by two carefully selected constrained mappings[4], which convert blocks of 16 address bits into blocks of 10nts, and blocks of 22 information bits into blocks of 13nts. Such design ensures that the maximum run length of G symbols is limited to three (to avoid G quadruplexes), and that the GC content is in the range of $40 - 60\%$.

#### 1.6 Encoding of Granular and Highly Relevant Image Features via LDPC Codes (Optional)

In some cases, fine facial details in pictures such as eyes and mouths may not be recovered properly due to our post blending processing. This leads to suboptimal reconstruction results, as shown in Supplementary Fig. 2, where the algorithm was unable to properly smooth out errors in the facial details such as lips and cheeks without blurring the images. Therefore, to further improve the reconstruction performance, a small amount of coding redundancy can be added to protect oligos that record selected facial features in eyes, noses, and lips so that they will not be smoothed out. Considering the error rate in practice and cost efficiency, we propose to use a regular, systematic low-density parity-check (LDPC) code to add the redundancy.

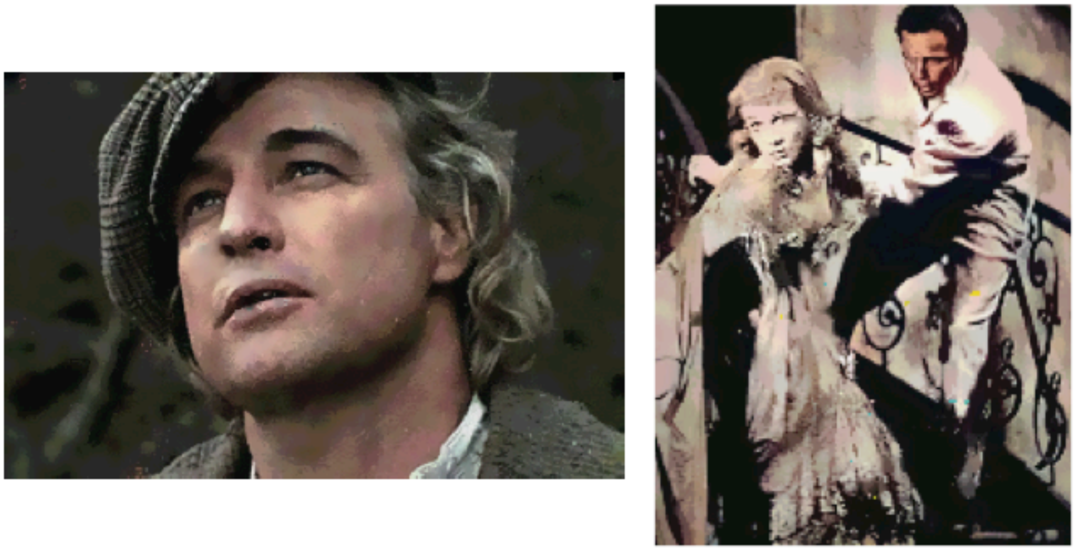

**Supplementary Figure 2.** Two examples showing that some facial details can be significantly blurred in the reconstructed images. The images in this figure are courtesy of: Paramount Pictures, Sony Pictures, MGM Studios, StudioCanal, American Zoetrope (© 1979 Zoetrope Corp. All Rights Reserved.), the Marlon Brando and Rod Steiger estates. The black and white public domain still of "A Streetcar Named Desire" was colorized using the software Hotpot.ai.

LDPC code[5] is a linear error correcting code, a method of transmitting a message over a noisy transmission channel, such as binary symmetric channel (BSC) and additive white Gaussian noise channel (AWGN). An LDPC is constructed using a sparse Tanner graph (subclass of the bipartite graph). LDPC codes are capacity-approaching codes, which means that practical constructions exist that allow the noise threshold to be set very close to the theoretical maximum (the Shannon limit) for a symmetric memoryless channel. The noise threshold defines an upper bound for the channel noise, up to which the probability of lost information can be made as small as desired. The maximum crossover probability LDPC code can correct decreases when code rate increases.

Using iterative belief propagation techniques, LDPC codes can be decoded in time linear to their block length.

In practice, the redundant information is encoded with a parity check matrix and decoded with a generator matrix. When systematic codes are used, information bits and parity check bits can be stored separately, which allows us to have a more compact design for oligos storing parity check bits (Fig. 1b in the main text). A parity check matrix is typically a sparse matrix, where the number of ones in a column $j$ (also the number of parity-check equations involving each code bit) and number of ones in a row $k$ (also the number of code bits involved in each parity-check equation) defines the code rate $R = 1 - j/k$. The length of a codeword is recorded as $n$. In our experiments, the $(n = 1200,\ j = 3,\ k = 12)$ systematic LDPC code with code rate 0.75 are used. The generated parity check bits are first concatenated to form a long binary string, then split into sub-blocks to be converted to DNA oligos of length 222nts following the data organization scheme shown in Fig. 1b. Compared to the organization scheme shown in Fig. 1a, there is no explicit address block, but the order of how to concatenate the binary strings read from DNA oligos is stored in the first 10bits. In addition to the parity-check bits, we also record information about oligos that represent facial details (and are subject to LDPC coding). The indices of facial feature oligos are encoded according to their address blocks. Bits are then converted to DNA oligos following the same organization in Fig. 1b. These oligos bearing indices are then repeated two times to form a 3-repition code, making the whole system more robust.

The new design only requires $265 + 42 \times 3 = 391$ more oligos, which is only of $391/11826 = 3.3\%$ redundancy compared to previous pool without using any error-correction redundancy. However, significant improvements in image quality can be easily observed in Fig. 2i.

Here, note that we use an unequal error-protection technique[6], which refers to adding highly limited redundancy only to oligos bearing facial features. Compared to conventional approaches[7,8] that add parity check for each part of the image, regardless of the content thereof, our approach uses much less memory and computational power. Thus, unequal error-protection in conjunction with our post-processing techniques enables significant reductions in costly error-correcting coding redundancy.

#### 2. Decoding Procedure for Sequence Dimension

The decoding procedure operates on the consensus reads from sequencing results and reverses the two-step encoding process. Note that when converting DNA consensus sequences back to binary strings, if one oligo unique identifier is corrupted by errors and its corresponding binary string does not exist, we will replace the erroneous identifier by another one at smallest Hamming distance from it. Each DNA block is able to be converted into some binary string, although this string may be wrong and cause visible discolorations in reconstructed image.

#### 3. Post-Processing for Sequence Dimension

To correct image discolorations, we implement a three-step postprocessing procedure. The first step is detecting the locations with discolorations, masking the regions out and subsequently treating them as missing pixels. The second step involves using deep learning techniques to inpaint the missing pixels. The third step involves smoothing the image to reduce both blocking effects caused by aggressive quantization and the mismatched inpainted pixels, and possibly including additional image enhancement features to further improve image quality for images with not granular faces.

##### 3.1 Automatic Discoloration Detection

Detecting arbitrarily shaped discolorations is a difficult problem in computer vision that has not been successfully addressed for classical image processing systems. This is due to the fact that discolored pixels usually have simultaneous distortions in all three-color channels of possibly different degrees. However, detecting discolorations in DNA-encoded images is possible since, with high probability, only one of the three-color channels at some positions will be corrupted due to independent encoding of the RGB components into different oligos. Thus, when two of the three channels are smooth in a particular region and the third channel is not smooth in that region (e.g., with pixel values that vary by more than a pre-determined extent), the variations in the third channel are likely due to error. We can then mask these regions out as discoloration regions. Another way to interpret our method is to view the RGB channels as a form of 3-repetition code, since one can use majority vote logic to determine which parts of the grayscale images are corrupted.

Supplementary Fig. 3a illustrates this fact that erroneous pixels in different channels do not overlap. The histograms in Supplementary Fig. 3b show that pixels with the smallest 15 frequencies values in the difference matrices $R - G, R - B, G - B$ indeed correspond to almost all erroneous regions in RGB channels, leaving out a few positions that are not covered though. Thus, we also perform an expansion of the identified regions to obtain the final masks in Supplementary Fig. 3c. Note that the masks, which are whitened out regions in the figures, are treated as missing data in images, and filled in using inpainting techniques in the next step.

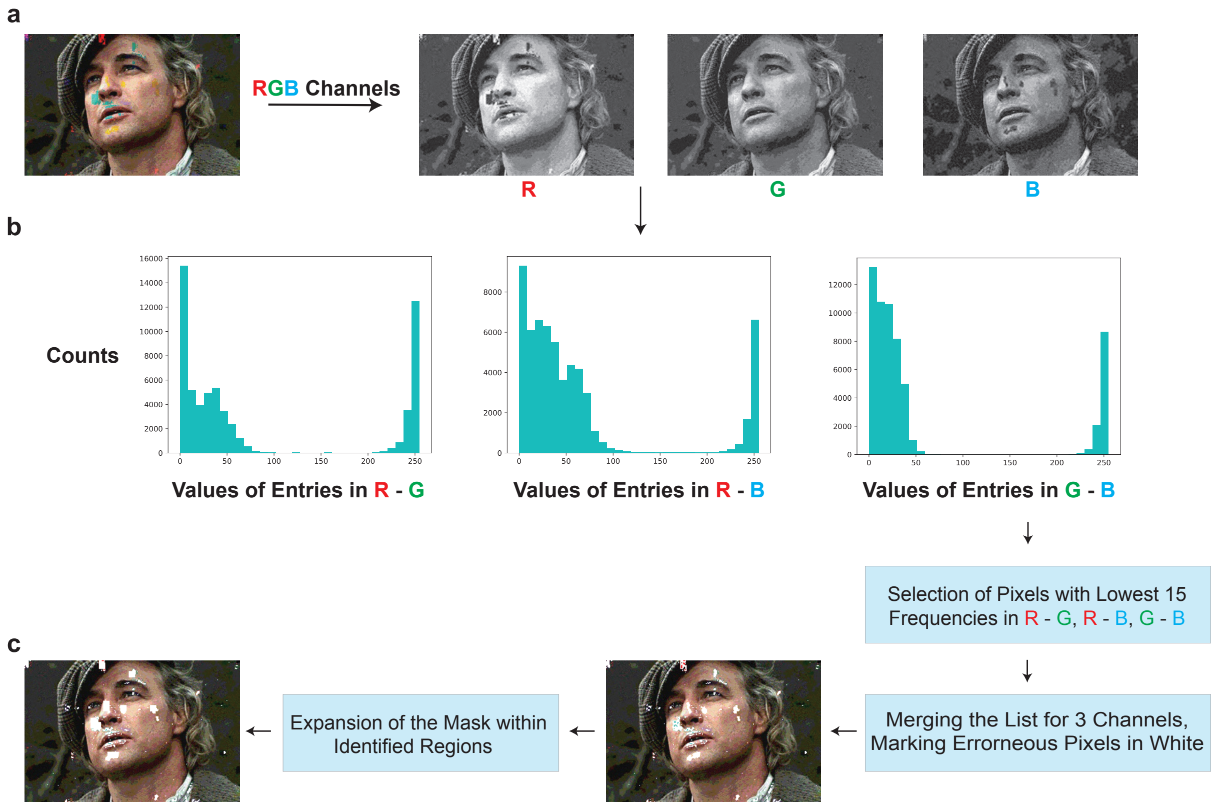


**Supplementary Figure 3.** Automatic discoloration detection scheme. The images in this figure are courtesy of: Paramount Pictures, Sony Pictures, MGM Studios, StudioCanal, American Zoetrope (© 1979 Zoetrope Corp. All Rights Reserved.), the Marlon Brando and Rod Steiger estates. Source data are provided as a Source Data file.

#### 3.2 Image Inpainting

Image inpainting, or image completion, is a method for filling out missing regions in an image. There exist several methods for image inpainting currently in use, including diffusion-based, patch-based[9] and deep learning approaches[10–12]. The former two methods use local or nonlocal information only within the target image itself which leads to poor performance when trying to recover complex details in large images. On the other hand, deep-learning methods such as GatedConvolution[11] and EdgeConnect[10] combine edges in the missing regions with color and texture information from the remainder of the image to fill in the missing pixels, as shown in Supplementary Fig. 4a. The architecture of one inpainting method that we use (EdgeConnect) is shown in Supplementary Fig. 4b. It consists of three steps: The first step involves extracting the edges of corrupted grayscale image via Canny edge detectors; the second step is to generate predicted edge maps that help with filling in the missing edges that are shown in red; the last step consists in generating the inpainting results based on the predicted edge map and corrupted color image.

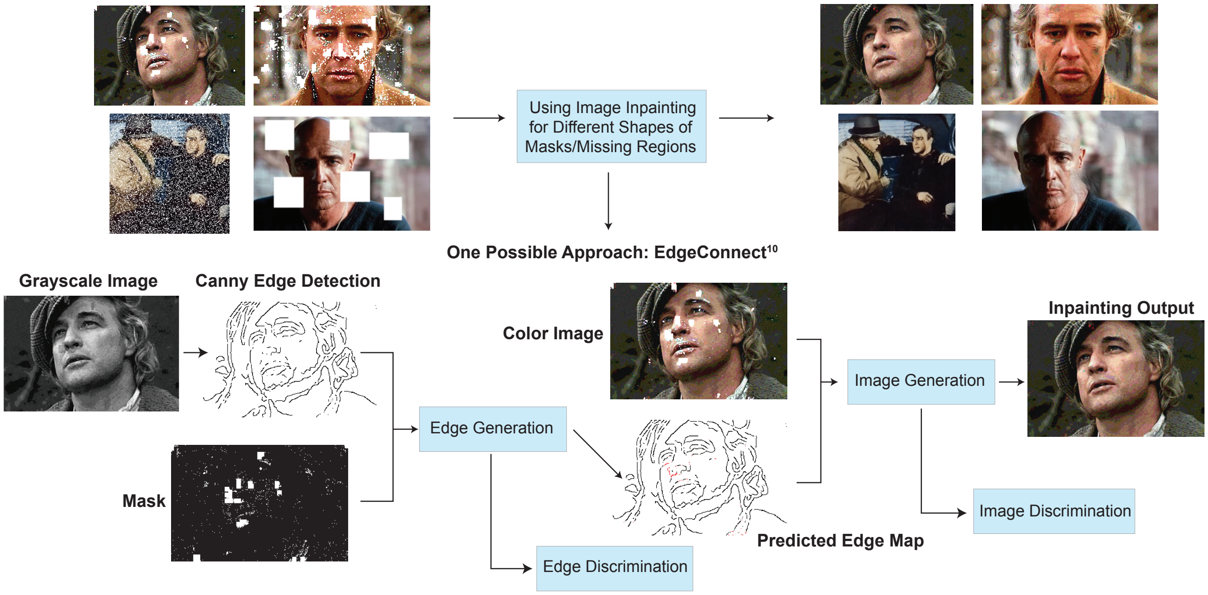


**Supplementary Figure 4.** Image inpainting scheme. The images in this figure are courtesy of: Paramount Pictures, Sony Pictures, MGM Studios, StudioCanal, American Zoetrope (© 1979 Zoetrope Corp. All Rights Reserved.), the Marlon Brando and Rod Steiger estates.

#### 3.3 Image Smoothing and Enhancement

Although the problem of discoloration may be addressed through inpainting, the reconstructed images can still suffer from mismatched inpainting and blocking effects caused by quantization. To further improve the image quality, smoothing is performed through bilateral filtering[15] that tends to preserve the edges structures. The smoothing equations read as:

$$\hat{I}[i,j] = \frac{\sum_{[k,l]\in\Omega} I[k,l]w\,(i,j,k,l)}{\sum_{[k,l]\in\Omega} w\,(i,j,k,l)},$$

$$w(i,j,k,l) = \exp\left(-\frac{(i-k)^2+(j-l)^2}{2\sigma_d^2} - \frac{\|I[i,j]-I[k,l]\|^2}{2\sigma_r^2}\right),$$

where $I$ denotes the original image and $\hat{I}$ the filtered image, $\Omega$ is some predefined window centered at the coordinates $[i,j]$, and $\sigma_d, \sigma_r$ are parameters that control the smoothing differences for intensities and coordinates, respectively. The filter performs Gaussian blurring on background regions but respects edge boundaries in the image. In our experiments, smoothing is performed with $\sigma_d^2 = \sigma_r^2 = 45$ and $\Omega$ of the form of a $9 \times 9$ square and no obvious discolorations are detectable. Furthermore, in order to address other possible impairments, the positions of error blocks, obtained from the discoloration detection platform, were used to perform adaptive median smoothing[13] around erroneous regions. As the final step, we may include additional image enhancement features[14] to further improve the quality of reconstructed images with no granular face details. The procedure is shown in Supplementary Fig. 5. Note that using image enhancement directly on the images generated from the information encoded in DNA oligos without error-correction redundancy results in poor image quality because classical image enhancement methods cannot automatically correct discoloration errors (Fig. 2k in the main text).

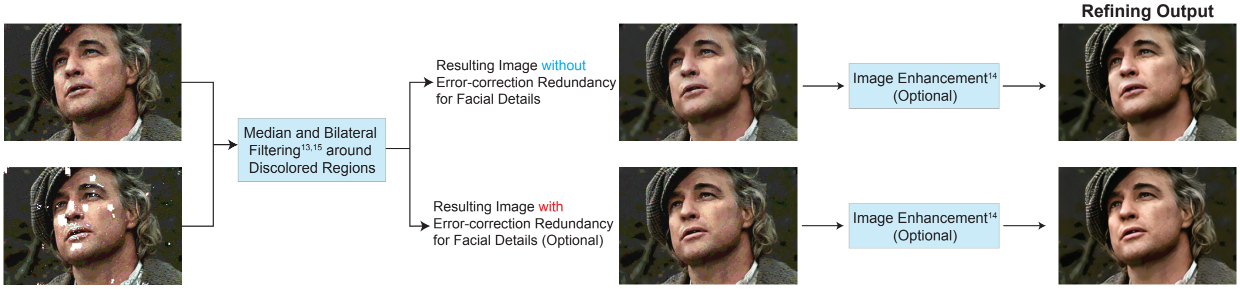


**Supplementary Figure 5.** Image smoothing and enhancement scheme. The images in this figure are courtesy of: Paramount Pictures, Sony Pictures, MGM Studios, StudioCanal, American Zoetrope (© 1979 Zoetrope Corp. All Rights Reserved.), the Marlon Brando and Rod Steiger estates.

### Supplementary Discussion

#### Additional Results on ML-Based Reconstruction versus JPEG with LDPC Coding Redundancy

As JPEG formats are highly sensitive to errors, even one mismatched nucleotide can result in poor-quality or unrecognizable image reconstruction (Supplementary Fig. 6). Hence, JPEG-encoded images need to be stored with coding overhead that guarantees the removal of all errors. Furthermore, even when coding overhead is present, and codes on graphs such as LDPC or turbo codes are used to generate the redundancy, one has to address the problem of channel parameter mismatch which can lead to additional issues with the reconstruction quality of the images.

To illustrate this point, we first perform simulated experiments in which we introduce one single substitution into the compressed JPEG file. In many cases, we observe obvious defects in the decoded image which cannot be fixed by current image processing techniques, as shown in Supplementary Fig. 6. In addition, if we by chance change the last bit in a JPEG compressed file, the JPEG decoding procedure terminates prematurely since the end-of-file identifier is missing.

To further illustrate the problems associated with the proposed source and channel coding schemes, we present an analysis pertaining to six combinations of JPEG image compression qualities and error-control coding schemes. For JPEG-compressed files with different quality parameters (recall that the quality parameter is a parameter in the Python Pillow Package for all image formats, JPEG included, that controls the quality of compressed images, and in on a scale from 1 (worst) to 95 (best)), we added low-density parity-check (LDPC) redundancy to the compressed data for error correction. For LDPC coding, we used both the standard approach and an approach known in the coding theory literature as interleaving (similar to shuffling[7]), which has the advantage of decorrelating the errors in oligos (Supplementary Fig. 8a, where we explained the interleaving process – writing blocks of user information row-wise, reading them out for encoding and recording on oligos column-wise). For all subsequent comparisons between our method and JPEG + LDPC, we made sure that the resulting oligo numbers (total file sizes) are matched as close as possible. For the standard approach, we encoded all binary information into DNA oligos and simulated a base substitution error rate of 0.8% and missing oligo error rate of 0.7%. These values match with what we observed in our real, synthetic oligos purchased from IDT, resulting in an overall bit error rate of 1.9%. We decoded the binary information from the erroneous DNA oligos using LDPC codes and followed decoding with JPEG reconstruction. The results are shown in Supplementary Fig. 7. Note that since JPEG has very specific formatting rules, missing or erroneous critical identifiers in JPEG files can lead to system errors, such as (OSError: broken data stream when reading image file) and (OSError: cannot identify image file) in Python. We included these error reports in our comparative analysis.

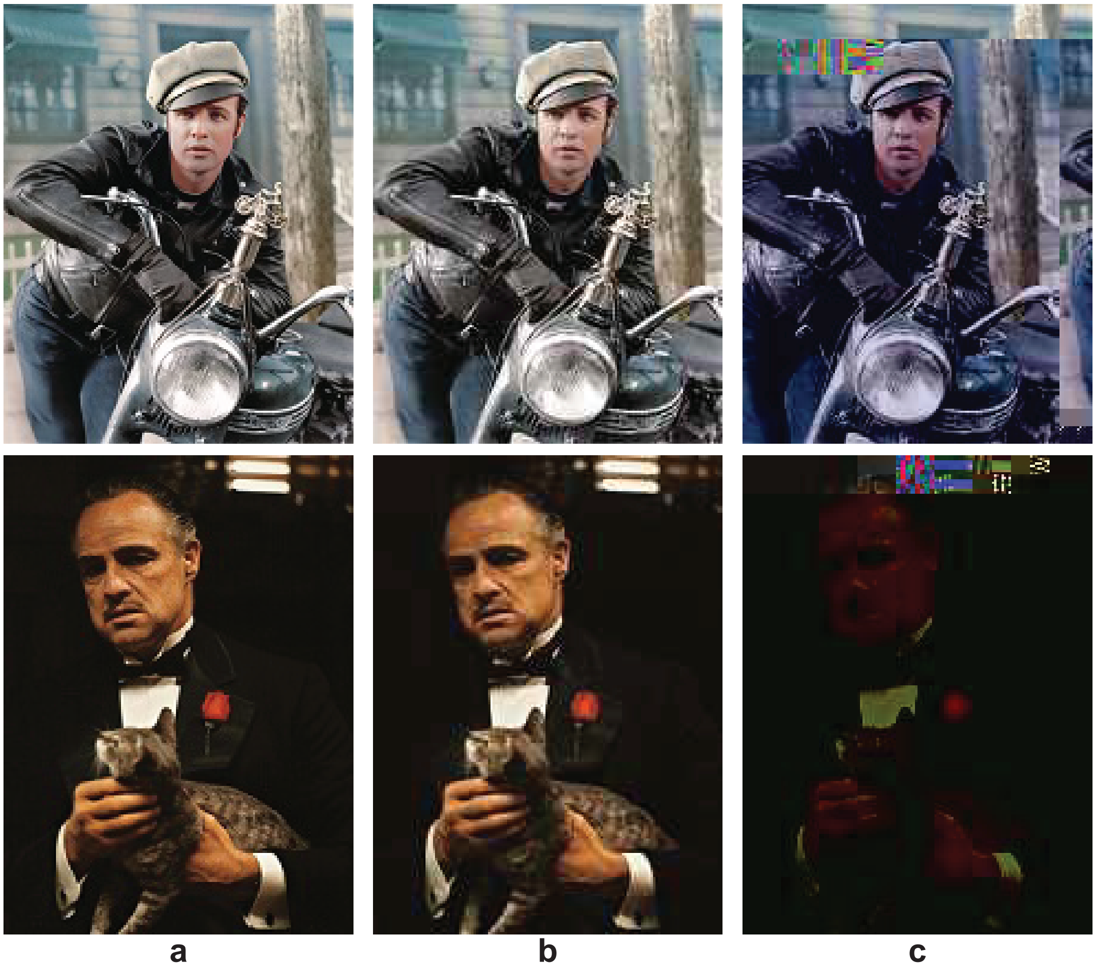


**Supplementary Figure 6.** Degradation in the quality of images containing decoding errors in the JPEG compressed files. **a)** The original image. **b)** The JPEG compressed image with quality parameter=40 (Top: PNSR=26.64, SSIM=0.91; Bottom: PNSR=30.25, SSIM=0.90). **c)** The results obtained after adding one substitution error in **b** (Top: PNSR=10.23, SSIM=0.10; Bottom: PNSR=13.60, SSIM=0.26). The images in this figure are courtesy of: Paramount Pictures, Sony Pictures, MGM Studios, StudioCanal, American Zoetrope (© 1979 Zoetrope Corp. All Rights Reserved.), the Marlon Brando and Rod Steiger estates. Source data are provided as a Source Data file.

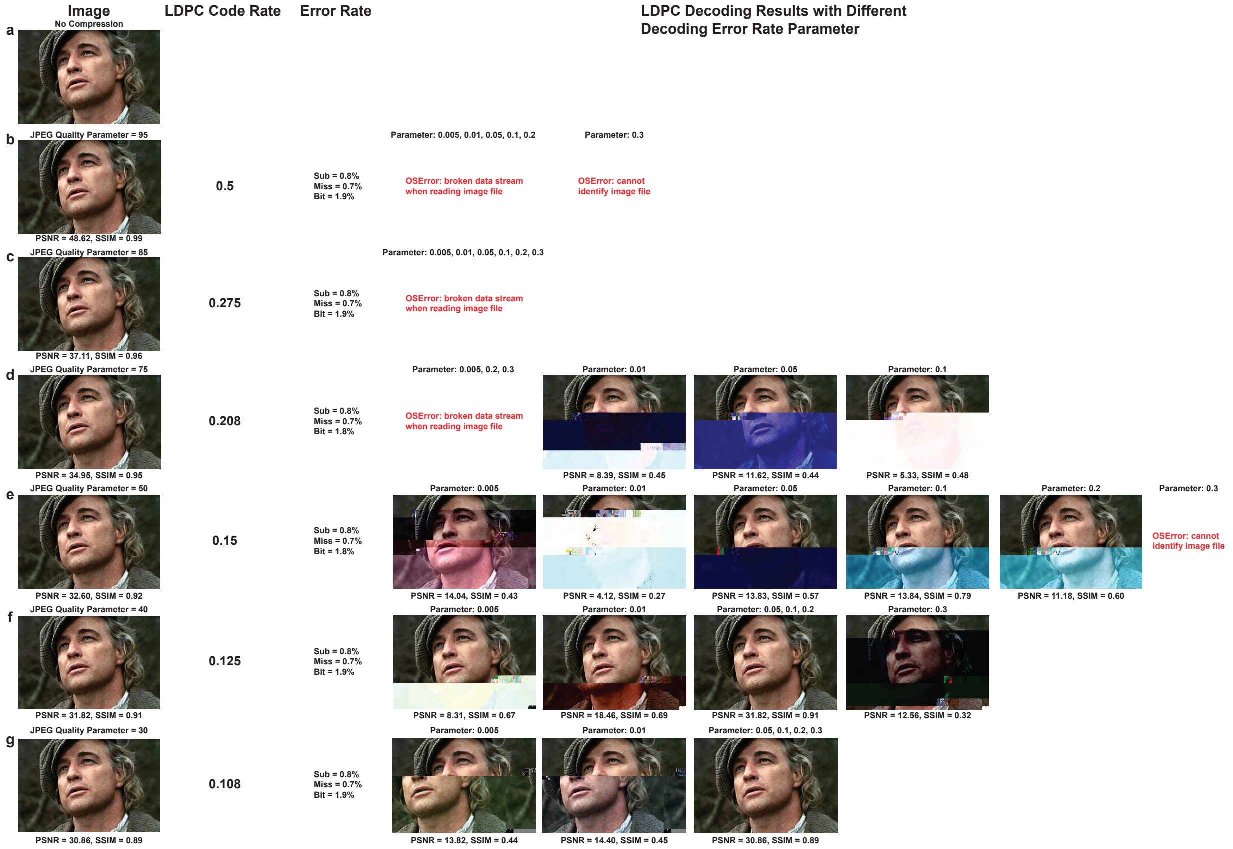


**Supplementary Figure 7**. Results of six combinations of JPEG image compression qualities and error-control coding schemes in the low error rate mode, when traditional LDPC coding is used. The three values in Column Error Rate are as follows: Substitution error (Sub), Missing oligo error (Miss) and overall bit error (Bit). We assume that the channel parameter for LDPC decoding is not known beforehand, as this parameter varies due to different sequencing quality, PCR errors, additional substitution errors arising due to rewriting etc. The images in this figure are courtesy of: Paramount Pictures, Sony Pictures, MGM Studios, StudioCanal, American Zoetrope (© 1979 Zoetrope Corp. All Rights Reserved.), the Marlon Brando and Rod Steiger estates. Source data are provided as a Source Data file.

Note that although the simulated bit error rate in some cases lies below the decoding "threshold" of the LDPC codes, the images may still not be correctly reconstructed as correlations in the errors may cause some oligos to be disproportionally affected and others to have barely any errors: For example, the missing oligo errors in the worst case can cause one whole LDPC codeword to be erased, which is then not possible to recover unless additional redundancy is added at the oligo levels. Oligo-level redundancy was used in the prior work by Grass et al.[16], but in our work it is replaced by interleaving followed by LDPC error-correction, as interleaving renders errors mostly uncorrelated and addresses the missing oligo issue without using Reed-Solomon codes. Even in this case, LDPC decoders may fails as the BSC transition probabilities are not known a priori, and mismatched decoding (i.e., decoding with the use of a wrong parameter estimate) is known to lead to errors. The results based on the proposed LDPC interleaving scheme are shown in Supplementary Fig. 8. Note that although interleaving can always provide us error-free decoding result when bit error rate is low, regardless of the estimated error probability set for decoding procedure (i.e., LDPC decoding can correct all bit errors when the base substitution error rate=0.8% and the missing oligo error rate=0.7%), it still cannot handle certain error scenarios. One case is when the error rate is actually higher than the one used for designing the codes (which arises, for example, if either more rewriting cycles are performed or the sequencer had higher error rates in subsequent runs); another is when the channel parameter is not properly known or estimated, in which case the decoder uses wrong initialization and hence wrong aggregation/message passing rules. As may be seen from Supplementary Fig. 8, error-free decoding can only be achieved when the estimated error probability is close to the real error rate (see the images for precise numerical values). Note that our image reconstruction results for low error rates (i.e., substitution error rate=0.8% and missing oligo error rate=0.7%) are available in the main text, and when the error rate is high, one can simply use a basic 3-bit quantization scheme instead of the more complicated scheme described in the main text. The reconstructed images after our ML-based post-processing are presented in Supplementary Fig. 9. The results show that 2DDNA is able to eliminate the need for worst-case coding redundancy and avoid problems with mismatched decoding parameters, while at the same time accommodate progressively degraded oligo qualities.

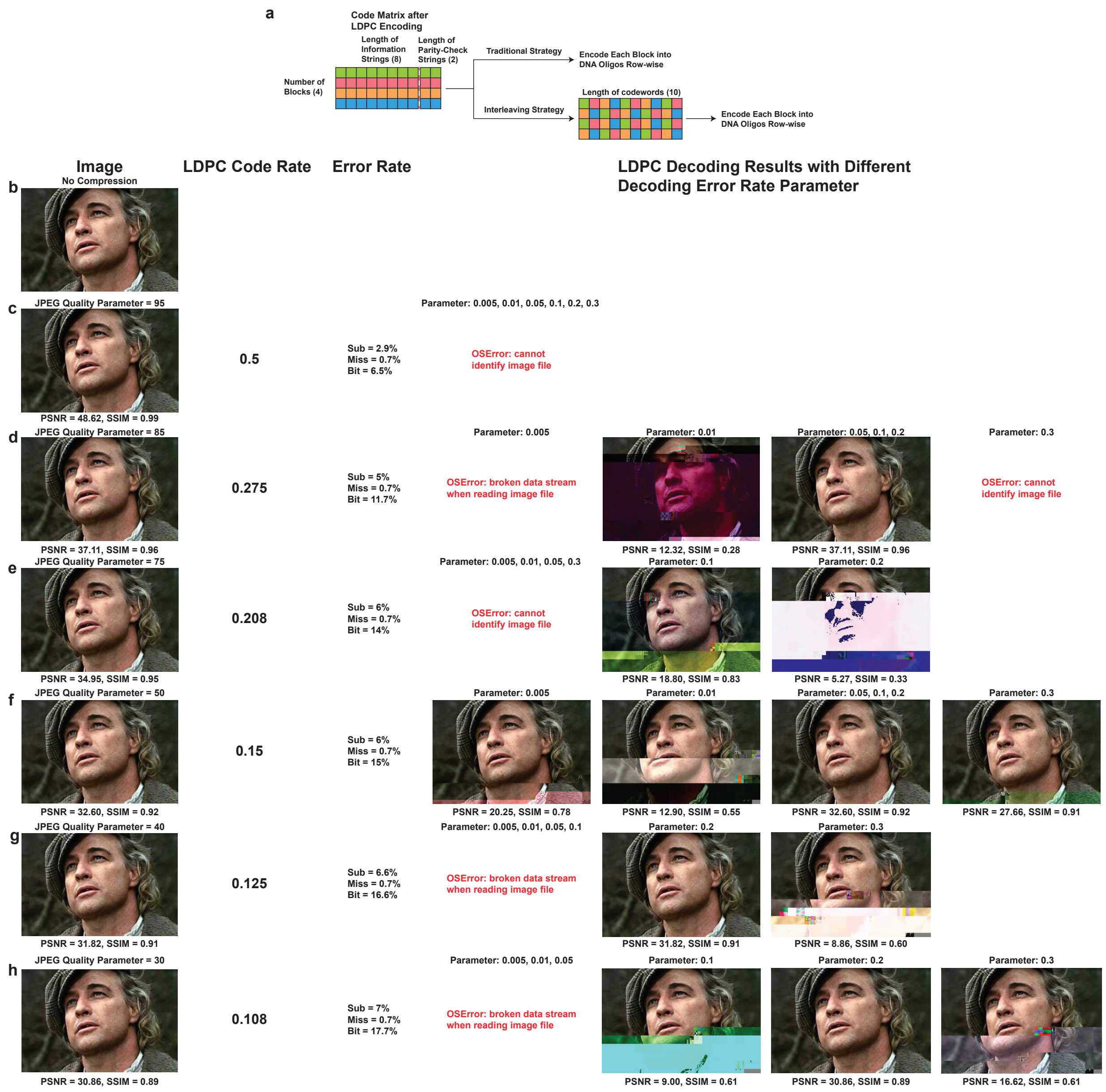


**Supplementary Figure 8**. **a**) Illustration of the traditional LDPC coding strategy and the interleaving strategy which aid with data recovery associated with missing oligos. In the traditional approach, we read the binary strings row-wise and encode them row-wise, while in the interleaving strategy, we read them row-wise and encode column-wise or vice-versa. This ensures that no block of consecutive pixels is missing and that the errors are more decorrelated. **b**) – **h**): Results of six combinations of JPEG image compression qualities and error-control coding schemes, for the case of high error rates and when interleaving is used. The images in this figure are courtesy of: Paramount Pictures, Sony Pictures, MGM Studios, StudioCanal, American Zoetrope (© 1979 Zoetrope Corp. All Rights Reserved.), the Marlon Brando and Rod Steiger estates. Source data are provided as a Source Data file.

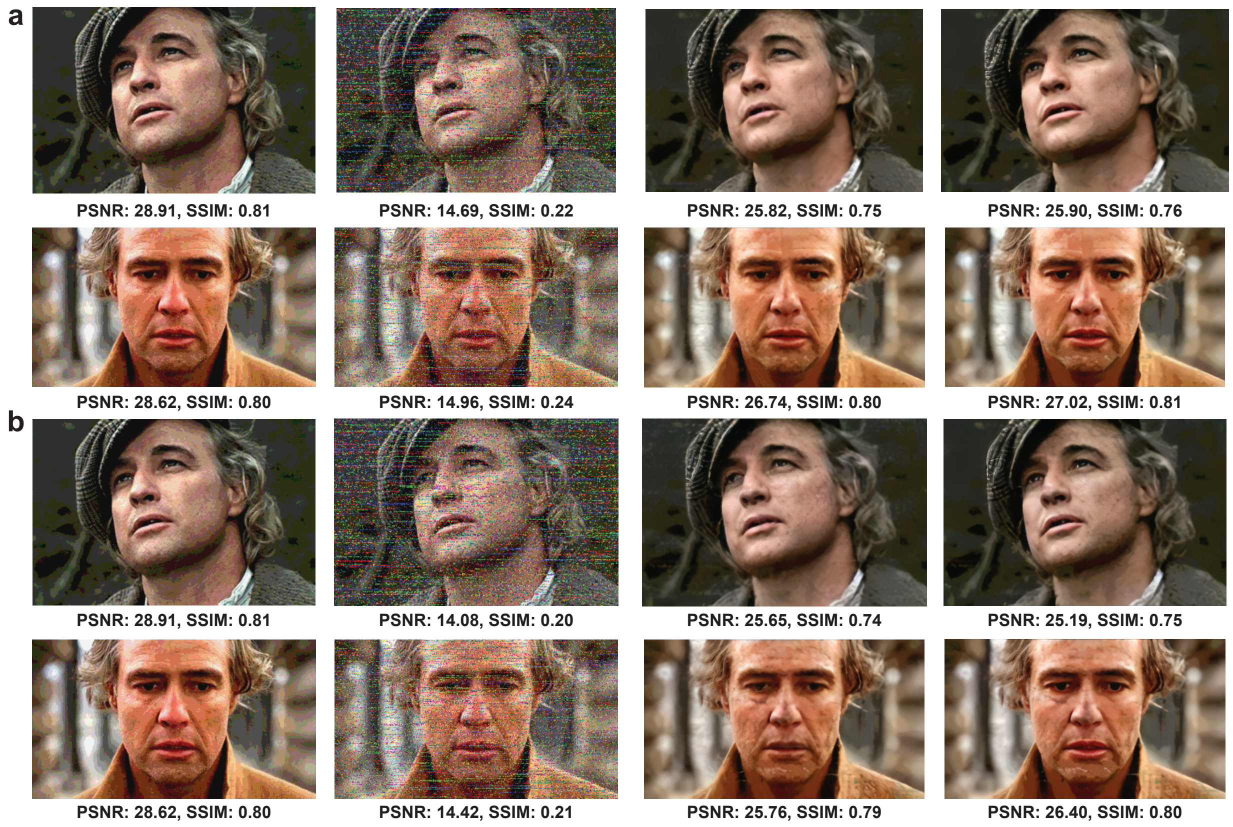


**Supplementary Figure 9**. **a**) The two images in the first column: 3-bit quantized versions of the original images, without lossless compression. The two images in the second column: Corrupted images decoded from DNA oligos with simulated errors (base substitution error rate = 6.6%, missing oligo error rate = 0.7%). The two images in the third column: Reconstructions obtained via our ML method without using unequal error protection redundancy. The two images in the fourth column: Reconstruction results based on our method with unequal error protection. **b**) The two images in the first column: 3-bit quantization of the original images. The two images in the second column: Corrupted images decoded from DNA oligos with simulated errors (base substitution error rate = 7%, missing oligo error rate = 0.7%). The two images in the third column: Reconstructions obtained using our ML-based method without using unequal error protection. The two images in the fourth column: Reconstructions obtained via our ML-based method with added unequal error protection redundancy. The images in this figure are courtesy of: Paramount Pictures, Sony Pictures, MGM Studios, StudioCanal, American Zoetrope (© 1979 Zoetrope Corp. All Rights Reserved.), the Marlon Brando and Rod Steiger estates. Source data are provided as a Source Data file.

#### Additional Results on JPEG versus GAN-Based Compression

Instead of JPEG, one may use a recently proposed compression approach based on GANs[17,18]. In this case, one needs to store not only the compressed image itself (the size of which is reported in terms of bits-per-pixel (bpp)), but also metadata describing how to decode the image with respect to the pre-trained models. In Supplementary Fig. 10, we provide a comparison of images compressed using JPEG with quality parameter 40 and the GAN-based method, with metrics and file sizes listed in the caption. Note that the GAN-based approach does not allow for arbitrary compression rates and only offers the user a choice between low, medium and high "quality". The image shown corresponds to medium quality and appears smoothed, and visually similarly to our image after processing. The file size for the medium quality setting is 1.2KB larger than that of the JPEG compressed image. Observe also that the JPEG compressed image is visually much more similar to the original image. We hence recommend using JPEG compressors, as these are not only standards and hence more practically relevant, but also better in terms of preserving image integrity under aggressive compression.

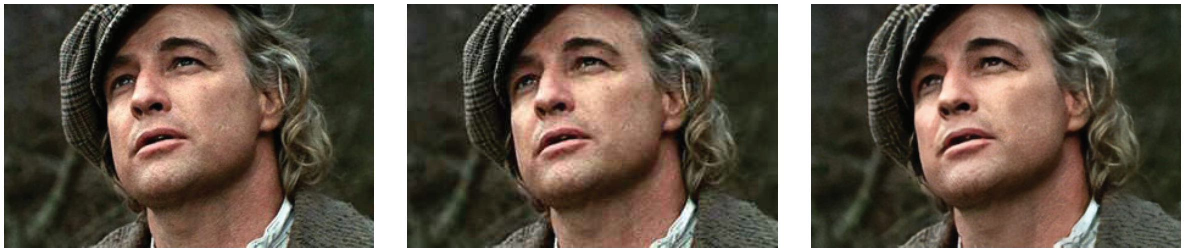

a b c

**Supplementary Figure 10**. **a**) The original, uncompressed image. **b**) The JPEG compressed image with quality parameter set to 40 (MSE=42.73, NRMSE=0.072, PSNR=31.82, SSIM=0.91, file size=6.8KB). **c**) The image compressed using the GAN-based approach from[17] (MSE=80.46, NRMSE=0.100, PSNR=29.07, SSIM=0.84, file size=8.0KB), which clearly involves a smoothing step that renders the resulting image less similar to the original but potentially more visually appealing. The images in this figure are courtesy of: Paramount Pictures, Sony Pictures, MGM Studios, StudioCanal, American Zoetrope (© 1979 Zoetrope Corp. All Rights Reserved.), the Marlon Brando and Rod Steiger estates. Source data are provided as a Source Data file.

#### Error Rate and Information Density

Before the first round of nicking, the number of mismatched nucleotides is 16,689nts, leading to an error rate of 0.0072 (0.72%). After nicking and ligation, the number of mismatched nucleotides goes up to 25,729nts, leading to an error rate of 0.0111 (1.11%).

In our experiments, a total number of 8,654,400bits was stored in 2,317,896nts, including address sequences, constrained balancing bits and markers. The information density of our platform equals the number of bits stored divided by the number of nucleotides used for encoding. Since quantization is used during the encoding procedure, there are two ways to compute this density: If calculated with respect to the number of bits in the raw image files, the information density equals 3.73bits/nt. Clearly, this exceeds the maximum 2 bits per nucleotide density dictated by the 4-alphabet size but may be seen as a consequence of the fact that we get a distorted image back, which allows for an increase from 2 to 3.73bits/nt. If the information density is calculated with respect to the number of bits of the quantized image files, the information density equals 1.40bits/nt. When converted into bytes/gram, the two reported densities theoretically equal 0.91 zettabytes/gram and 0.34 zettabytes/gram.

#### Supplementary Tables

Nine pairs of primer sequences used in our experiments are at a Hamming distance larger than 10nts and paired up to have similar melting temperature, which allows for all oligos to be amplified in the same cycle. The specific primer sequences along with their physical properties are shown in Supplementary Table 1.

| Prefix Primer | Melting Temperature | GC Content | Suffix Primer | Melting Temperature | GC Content |
|---|---|---|---|---|---|
| 5'-CCTTAGAAGT-CGCAATAAGT-3' | 49.6° C | 40% | 5'-AATTACTAAG-CGACCTCGTC-3' | 51.8° C | 45% |
| 5'-CAGATCGTCA-GGCCTATTAT-3' | 51.3° C | 45% | 5'-ATAACACGTG-TCGCTGTATC-3' | 52.4° C | 45% |
| 5'-GAGACGACCT-TTACACACTT-3' | 52° C | 45% | 5'-AATTCACGGT-CAGAGACATG-3' | 52.4° C | 45% |
| 5'-GTAACAAGTA-GGGTTGAGCA-3' | 52.1° C | 45% | 5'-TATACGCCTT-CAACGATGAG-3' | 51.9° C | 45% |
| 5'-GTCATCGAGT-CGATAGGTAT-3' | 50.6° C | 45% | 5'-AACACATGTC-ATCGAGTCTC-3' | 52.1° C | 45% |
| 5'-GAGAGCGAGT-AGAGTACAAA-3' | 51° C | 45% | 5'-TGTAGCAGTC-ACTTCTTCTC-3' | 51.5° C | 45% |
| 5'-GGGTGGTTAG-AAGTTGTATT-3' | 49.6° C | 40% | 5'-AGTGAGTCAT-CCTAGTTCTC-3' | 50.5° C | 45% |
| 5'-GGACTGCCTG-GAACTATTAA-3' | 51.7° C | 45% | 5'-ATACACTCAT-AACACCTCGG-3' | 51.4° C | 45% |
| 5'-GGACATATAT-GTGCTGACCA-3' | 51.8° C | 45% | 5'-GAGTCCAGTA-TGAATCTCGT-3' | 51.1° C | 45% |

**Supplementary Table 1.** Nine pairs of primer sequences used in our experiments.

| Enzyme | Recognition Sequence | Buffer | | Temperature |
|---|---|---|---|---|
| | | CutSmart | NEBuffer 3.1 | |
| Nb.BtsI | 5’-GCAGTGNN-3’<br>3’-CGTCAC\|NN-5’ | ✖ | | 37° C |
| Nt.BstNBI | 5’-GAGTC\|NNNNN-3’<br>3’-CTCAGNNNNN-5’ | | ✖ | 55° C |
| Nb.BssSI | 5’-CACGAG-3’<br>3’-GTGCT\|C-3’ | | ✖ | 37° C |
| Nt.AlwI | 5’-GGATCNNNN\|N-3’<br>3’-CCTAGNNNNN-5’ | ✖ | | 37° C |
| Nt.BsmAI | 5’-GTCTCN\|N-3’<br>3’-CAGAGNN-5 | ✖ | | 37° C |
| Nb.BsmI | 5’-GAATGCN-3’<br>3’-CTTAC\|GN-3’ | | ✖ | 65° C |
| Nb.BsrDI | 5’-GCAATGNN-3’<br>3’-CGTTAC\|NN-5’ | ✖ | ✖ | 37° C |
| Nb.BbvCI | 5’-CCTCAGC-3’<br>3’-GGAGT\|CG-5’ | ✖ | | 37° C |
| Nt.BbvCI | 5’-CC\|TCAGC-3’<br>3’-GGAGTCG-5’ | ✖ | | 37° C |
| Nt.BspQI | 5’-GCTCTTCN\|-3’<br>3’-CGAGAAGN-5’ | | ✖ | 50° C |

**Supplementary Table 2.** Enzymes used in the nicking experiments (writing and rewriting).
